# Glucose selectively drives a rapid oxidative burst and immunometabolic reprogramming in human neutrophils during *Mycobacterium tuberculosis* infection

**DOI:** 10.64898/2026.05.05.722986

**Authors:** Krishna C. Chinta, Delon Naicker, Sajid Nadeem, Ritesh Sevalkar, Vanessa Pillay, Threnesan Naidoo, Gordon Wells, Kapongo Lumamba, Kievershen Nargan, Ashendree Govender, Jeff J. Jones, Marion Pang, Baiyi Quan, Ting-Yu Wang, Michael L. Roukes, Dongquan Chen, Hayden T. Pacl, Anupam Agarwal, Joel N. Glasgow, Adrie J. C. Steyn

## Abstract

Neutrophil functions have been linked to tuberculosis (TB)-associated tissue damage; however, the mechanisms driving immunopathology in the human TB lung remain poorly understood, due partly to the scarcity of human tissue for study. Here, we examine the metabolic and bioenergetic reprogramming of human neutrophils in response to *Mycobacterium tuberculosis* (*Mtb*) infection. In human necrotic TB granulomas, levels of NETosis-associated proteins are increased and co-localize with GLUT3, linking nutrient uptake to tissue damage. *In vitro*, *Mtb* elicits an immediate, contact-dependent oxidative burst in human neutrophils, and the magnitude of this response is carbon source-dependent. Glucose enables the most robust responses, indicating that glucose metabolism is a key driver of neutrophil-mediated inflammatory damage during TB. *Mtb*-induced responses are distinct from those induced by PMA, non-tuberculous mycobacteria, or other pathogenic intracellular bacteria, and are mediated through multiple neutrophil surface receptors. Notably, our data show that while the oxidative burst is carbon source-dependent, cytokine production is not. Further, *Mtb* infection reprograms neutrophil metabolism from glycolysis to the pentose phosphate pathway (PPP), generating NADPH required for the oxidative burst. Inhibiting G6PD, NADPH oxidase, or PAD4 significantly reduces this response, highlighting the PPP as a promising host target for mitigating TB immunopathology.

## Introduction

Tuberculosis (TB), caused by *Mycobacterium tuberculosis* (*Mtb*), is the leading cause of human mortality from a single infectious agent^1^. Major clinical manifestations of pulmonary TB include reduced lung function and shortness of breath, hemorrhage, dysregulated immune responses, and significant tissue damage that often requires resection in patients with active TB^2–6^. Consequently, there is an urgent need to develop host-directed therapies (HDTs) capable of modulating inflammatory responses to reduce TB-associated immunopathology^7,8^. However, there are significant gaps in our understanding of how host factors drive dysregulated immune responses and tissue pathology during TB^9^.

During *Mtb* infection, host immune responses lead to the formation of granulomas, organized structures consisting mainly of myeloid cells and lymphocytes, which act to restrict bacterial spread^10^. Within the myeloid compartment, neutrophils are often elevated in the bronchoalveolar lavage fluid, sputum, and necrotic lesions of patients with TB, and neutrophil abundance correlates with granuloma necrosis and poor disease outcomes^11,12^. Neutrophils isolated from severely damaged regions of human TB lungs exhibit elevated levels of reactive oxygen and nitrogen species (ROS and RNS), suggesting that failure to regulate free radical responses contributes to host tissue damage^4^. Yet, the mechanisms that underly neutrophil-driven immunopathology in TB are incompletely understood. Moreover, it is still not clear whether the granuloma microenvironment itself can influence the effector functions of neutrophils or other immune cells^13^.

In mouse models of TB, depleting neutrophils with Ly6G antibodies reduces bacterial burden and TB-associated immunopathology^14–16^. However, the translational potential of this approach is limited, since depletion of neutrophils could severely compromise innate immune defenses and increase susceptibility to various secondary infections^17,18^. Rather, selective inhibition of neutrophil functions that contribute to tissue pathology would represent a more viable approach. For example, interventions that limit neutrophil extracellular trap formation (NETosis), a major driver of tissue necrosis in animal models of TB^14,19^, could provide therapeutic benefit. In *Mtb*-infected mice, administration of the pan–protein arginine deiminase (PAD) inhibitor chloro-amidine (Cl-amidine) or the selective PAD4 inhibitor GSK-484 significantly reduces NETosis, necrosis, tissue damage, and bacillary burden^19^. These findings highlight the therapeutic potential of targeting pathogenic neutrophil functions as an adjunctive HDT component of TB treatment regimens.

The TB granuloma microenvironment shares certain immunological characteristics with that of solid tumors, including its immune cell composition, chronic, dysregulated inflammation, TGF-β release, and mechanisms of immunosuppression^20,21^. Tumors can create an immunosuppressive local milieu that hinders effective immune responses; this is achieved in part by increasing glucose uptake and metabolism, resulting in a nutrient-deprived microenvironment that is detrimental to immune effector functions^22,23^ . While the metabolism-driven immunomodulatory effects of tumors have been well studied, a comprehensive characterization of the human TB granuloma microenvironment and its impact on host immunometabolism is still lacking. *Mtb* induces a metabolically quiescent state in human macrophages *in vitro* and in CD8⁺ T cells from infected mice, enforcing metabolic rewiring and altering effector responses^24,25^. In mouse and human macrophages, *Mtb* infection suppresses glycolysis and oxidative phosphorylation (OXPHOS)^24^. Furthermore, *Mtb* infection of macrophages results in NAD⁺ depletion, thereby suppressing glycolysis and impairing host-protective IFN-γ responses^26,27^. Similarly, *Mtb* infection in mice shifts CD8⁺ T-cell metabolism from predominantly OXPHOS toward glycolysis, leading to upregulation of inhibitory receptors and deterioration of CD8⁺ T-cell effector functions^25^. Importantly, treatment of *Mtb*-infected mice with nicotinamide^26^ or metformin^25^ restores glycolysis in macrophages and mitochondrial metabolism in CD8⁺ T cells, and suggests that targeting distinct steps in host cell metabolism could be a viable HDT approach against TB. Neutrophils are reliant on glycolysis to meet energy demands for their effector functions, including the oxidative burst and NETosis^28^, processes that correlate with increased expression of glucose transporter 1 (GLUT-1)^29,30^. However, upon chemical stimulation with phorbol-12-myristate-13-acetate (PMA) or in pathological conditions such as COPD, neutrophils can shift their metabolism toward the pentose phosphate pathway (PPP) or gluconeogenesis, switching to alternative carbon sources to sustain the energy demand for their effector responses^31,32^. Neutrophil immunometabolism in general is less well characterized than that of macrophages and T cells, largely due to neutrophils’ short lifespan and sensitivity to *ex vivo* manipulation^33,34^. A poor understanding of neutrophil immunometabolism has limited our ability to accurately describe the role of neutrophils in TB pathogenesis.

A significant barrier to defining the contributions of neutrophils toward human TB immunopathology is a heavy reliance on mouse models, which do not reproduce the structural and pathological complexity of human TB lesions^10,35,36^. Three-dimensional imaging of resected human TB lungs has revealed that TB granulomas are remarkably heterogeneous in size and morphology and are likely shaped by the bronchi, highlighting fundamental differences from mouse pathology^5,6^. Such complexities raise several critical questions. Does necrosis in human pulmonary granulomas correlate with NETosis, and is this relationship dependent solely on neutrophil glucose metabolism? What factors govern the transition from protective neutrophil activity, including anti-*Mtb* functions, to pathological responses that drive tissue damage? Can we identify distinct metabolic signatures in neutrophils that predict disease trajectory or outcomes? Finally, can neutrophil metabolism be pharmacologically modulated to limit lung destruction in patients with TB without compromising essential antimicrobial defenses?

In this study, we hypothesized that *Mtb* infection reprograms immunometabolism in human neutrophils leading to enhanced oxidative burst, ROS production, and NETosis. To test this hypothesis, we used proteomic and multiplex immunofluorescence (mIF) analyses to characterize the expression patterns of glucose transporters and proteins associated with neutrophil effector functions in necrotic human TB lung lesions. We then performed *in vivo* infection assays using neutrophils isolated from healthy donors to assess how *Mtb* infection and glucose availability influence neutrophil viability, oxidative burst, ROS production, and NETosis. Using metabolomic and real-time bioenergetic analyses, we measured *Mtb*-induced changes in neutrophil metabolism. Further, inhibition of glucose-6-phosphate dehydrogenase (G6PD), NADPH oxidase (NOX2) and PAD4 identified the metabolic pathways driving these responses. Our data demonstrate that *Mtb* infection and glucose availability are critical in shaping neutrophil oxidative burst and metabolic reprogramming, and highlight the PPP, glucose transporters, and NETosis as potential therapeutic targets for limiting TB-driven immunopathology.

## Results

### Proteomic profiling shows enrichment of NETosis-associated proteins in human necrotic granulomas

Neutrophils are increasingly recognized as key mediators of TB immunopathology; however, animal models may not accurately reproduce the neutrophil effector functions, including NET formation, that occur in human TB^37^. For example, when compared to mouse neutrophils, the total protein mass of human peripheral neutrophils is significantly higher, with human neutrophils containing significantly more primary granule proteins, including myeloperoxidase (MPO), effector proteases, and NADPH oxidase (NOX)^37^. While a few human neutrophil studies exist^38,39^, the contribution of NET components such as citrullinated histones, NE, and MPO to the development of tissue necrosis in human TB has been overlooked.

Caseous necrotic granulomas are a key characteristic of chronic human pulmonary TB, yet the molecular and cellular factors that drive TB immunopathology in these lesions remain poorly understood, highlighting the need for high-resolution analyses. In this regard, proteomic analysis of human TB granulomas revealed enrichment of pro- and anti-inflammatory proteins^40^. Nonetheless, critical gaps remain in our understanding of how neutrophil-driven processes, especially NETosis, may contribute to the immunopathology characteristic of necrotic human TB granulomas. To address this, we first performed laser capture microdissection (LCM) to isolate a single necrotic granulomatous region from each of 27 well-characterized human TB lung specimens obtained from 17 patients with TB who had undergone lung resection, as well as 6 healthy lung specimens from 2 patients who died of non-TB causes (Fig. 1a, Extended Data Table. 1). Liquid chromatography–mass spectrometry (LC-MS) followed by proteomic analysis identified 4,772 proteins in necrotic granulomatous specimens, and the abundance of 1,228 proteins was significantly different compared to healthy lung tissue from patients without TB (Fig. 1a,b). Of these 1,228 proteins, 173 were neutrophil-associated proteins, with 135 more abundant and 36 less abundant in necrotic TB lesions compared to healthy lung tissue (Fig. 1b, Extended Data Table 2). Notably, we observed consistent enrichment of 51 NETosis-associated proteins in each of the 27 necrotic LCM samples (Fig. 1c), with statistically significant increases in the abundance of 14 key NETosis-related proteins (Fig. 1d). These include the NETosis-inducing enzyme PAD4, core NET components azurocidin (AZU1), neutrophil elastase (ELANE/NE), and MPO, as well as CYBA/p22^phox^ and CYBB/NOX2/gp91^phox^, the essential membrane-bound subunits of the phagocyte NADPH oxidase 2 complex^41^ that generates ROS and the oxidative burst during NETosis. Further, the levels of NET-associated histone proteins H2BC12L, H2BC26, and H4C16 were significantly increased in necrotic regions, along with enzymes associated with glucose metabolism including hexokinase 3 (HK3), phosphofructokinase (PFKL), and pyruvate kinase (PKM) (Fig. 1d). The abundance of the high-affinity glucose transporter 3 (GLUT3) was significantly higher in necrotic granulomas than in healthy lung tissue, whereas GLUT1 was less abundant in necrotic lesions, suggesting a role for GLUT3 in driving neutrophil-mediated tissue necrosis in TB granulomas (Fig. 1d).

**Fig. 1.**
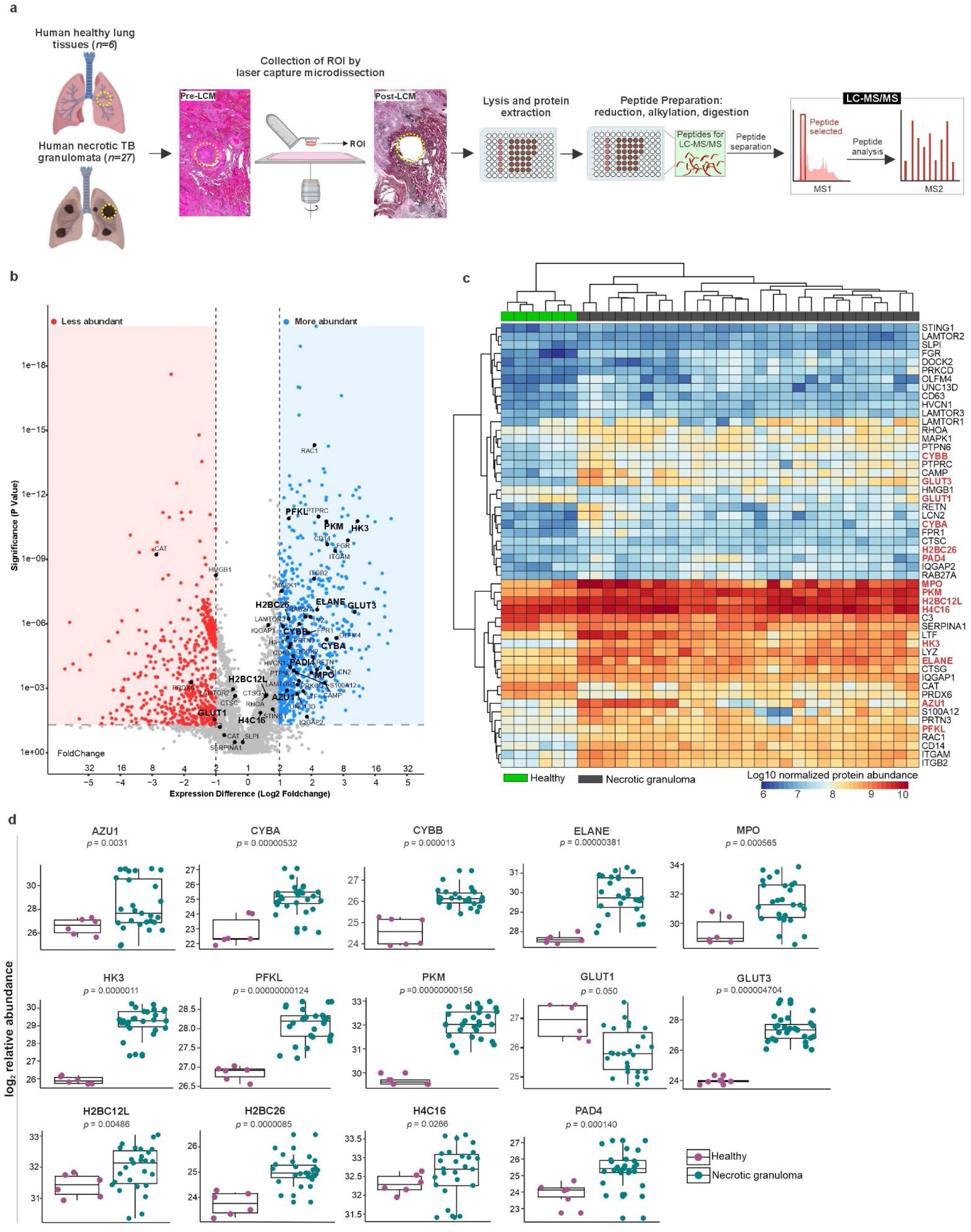
| **NETosis-associated proteins are enriched in human necrotic TB granulomas. a,** Workflow for isolation and proteomic analysis of human lung tissue specimens. Regions within 27 necrotic pulmonary TB granulomas obtained from 17 patients with TB, as well as regions of healthy lung tissue from two subjects who died of causes other than TB, were isolated by laser-capture microdissection (LCM) and subjected to LC–MS/MS-based proteomic analyses. **b**, Proteomic profiling of necrotic granulomas identified 4,772 proteins, each represented as a dot in the volcano plot. In necrotic granulomas, the abundance of 1,228 proteins was significantly higher (blue dots) or lower (red dots) compared to healthy control lung tissue. Proteins associated with neutrophil functions are in bold text. **c**, Hierarchical clustering of the log10 normalized protein abundances of 51 NETosis-associated proteins in 27 necrotic regions compared to healthy lung tissues. **d**, Relative abundance of key NETosis-related proteins in human necrotic granuloma and control tissues including PAD4, MPO, NE, azurocidin (AZU1), CYBA and CYBB, histone proteins (H2BC12L, H2BC26, H4C16), glycolytic enzymes (HK3, PFKL, PKL), GLUT1, GLUT3. Statistical significance was determined using unpaired, non-parametric Mann–Whitney test and the respective *P* values are indicated.

The co-enrichment of GLUT3, glycolysis enzymes, and NETosis-associated proteins in necrotic granulomas suggested that glucose availability differs between necrotic granulomatous and normal tissue. To investigate this, we compared the glucose content in human necrotic TB granulomas and non-TB lung control specimens (Extended Data Table 3). We observed no significant difference in glucose content between these groups (Extended Data Fig. 1a,b), suggesting that it is not glucose availability but rather glucose uptake and utilization, potentially mediated by GLUT3, that drives NETosis-associated necrosis in the TB granuloma.

In summary, proteomic analysis of isolated human necrotic granulomas reveals a consistent and robust enrichment of NETosis-associated proteins and key metabolic enzymes, particularly those involved in glucose metabolism. Our findings suggest that GLUT3-mediated glucose uptake, rather than glucose abundance *per se*, fuels neutrophil activation and NETosis-driven tissue damage. Further, elevated levels of core NET components, NETosis inducers, and glucose metabolism enzymes in necrotic granulomas provide compelling evidence for a link between neutrophil activation, metabolic reprogramming, and tissue necrosis in pulmonary TB.

### GLUT3 is spatially associated with NETosis in human necrotic TB granulomas

We next performed multiplex immunofluorescence (mIF) microscopy to validate our proteomic findings and determine the spatial distribution of GLUT1, GLUT3, and key NETosis markers MPO, NE, and citrullinated histone H3 (H3-Cit) within human necrotic granulomas. Consistent with studies that found NETosis markers in circulating neutrophils of patients with TB and caseating TB granulomas in non-human primates (NHPs)^19,42^, we detected NETosis-associated proteins in human necrotic granulomas (Fig. 2, Extended Data Fig. 2).

**Fig. 2.**
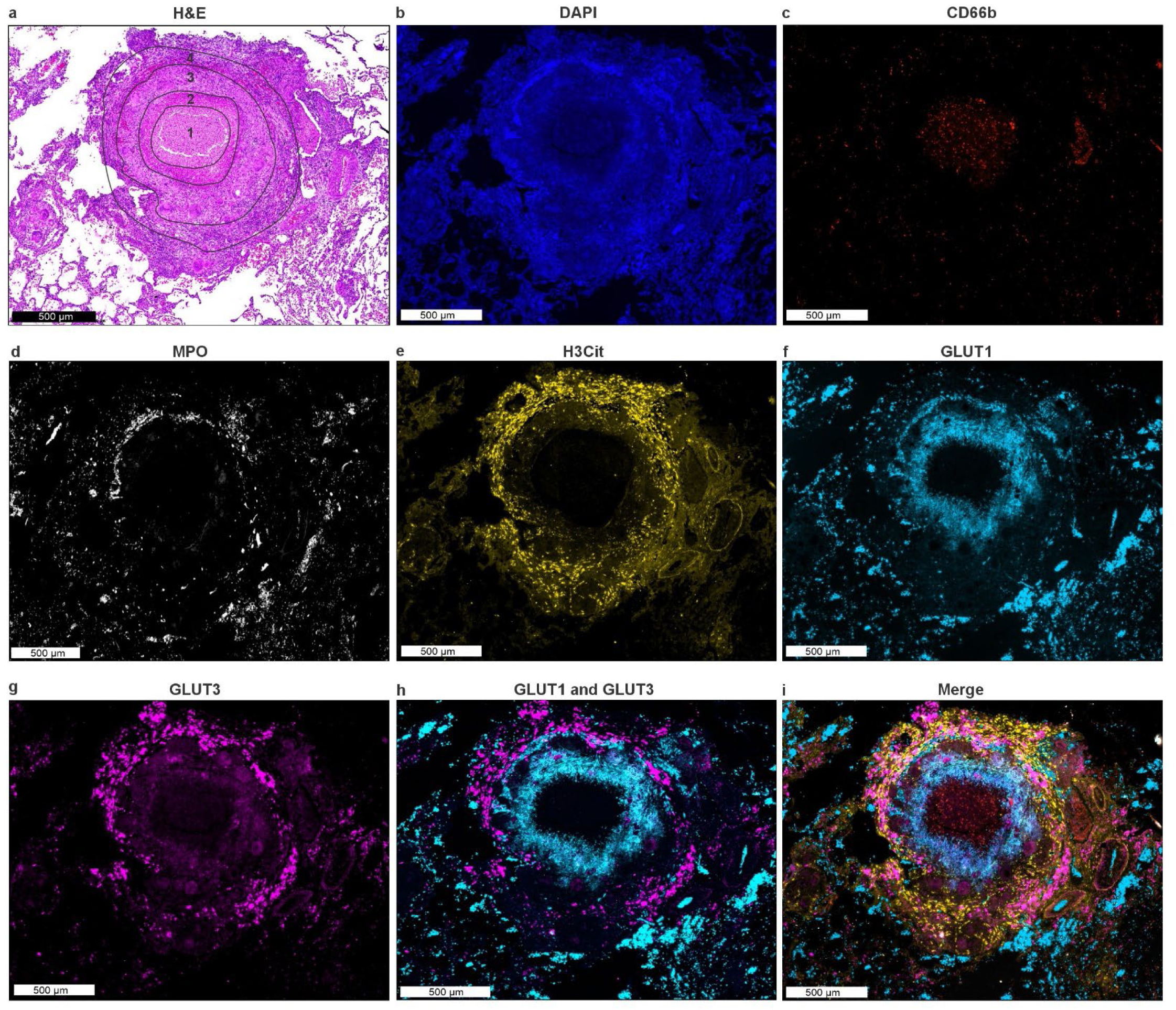
| **GLUT3 colocalizes with NETosis proteins in human necrotic TB granulomas**. Representative necrotic TB granuloma displaying four distinct pathological regions: (1) necrotic center, (2) circumference of necrotic center, (3) cellular periphery consisting of fibroblasts, macrophages, Langhans-type giant cells, and (4) interface between granuloma and adjacent tissue. H&E and DAPI staining and mIF microscopy were performed on a single tissue section. **a**, H&E staining of a representative necrotic TB granuloma. **b**, DAPI nuclear staining of the granuloma in (**a**). **c**-**i**, multiplex immunofluorescence (mIF) microscopy of the same tissue section revealing that CD66b^+^ neutrophils (**c**) colocalize with NETosis markers NE (**d**), MPO (**e**), and H3-Cit (**f**), particularly at the granuloma-tissue interface (region 4). **g**, GLUT1 is detected in regions 2 and 3 adjacent to the necrotic center where neutrophils and NETosis markers are scarce. **h,** GLUT3 is localized at the granuloma-tissue interface (region 4) and colocalizes with NETosis markers NE, MPO, and H3-Cit. **i**, merged mIF signals. Autofluorescence subtraction and image processing were performed using HORIZON software. Additional images of NETosis markers in necrotic granulomas are provided in Extended Data Fig. 2.

Necrotic TB granulomas were identified in human TB lung specimens and pathologically distinct regions within a representative granuloma were annotated as follows: 1. necrotic center, 2. outer rim of the necrotic center, 3. cellular periphery, 4. interface between granuloma and adjacent tissue (Fig. 2a). All H&E staining (Fig. 2a, Extended Data Fig. 2), DAPI staining (Fig. 2b, Extended Data Fig. 2), and mIF microscopy (Fig. 2b-h, Extended Data Fig. 2) were performed on single tissue sections to allow precise spatial registration of antigen and nuclear staining with lesion structure. Neutrophils were detected by CD66b expression (Fig. 2c) and were abundant within the necrotic granuloma, with the strongest signals observed in the necrotic core (Fig. 2a, region 1) and at the intersection of the granuloma and adjacent tissue (region 4). Notably, CD66b^+^ neutrophils at this intersection colocalize with NETosis markers MPO (Fig. 2d, Extended Data Fig. 2) and H3-Cit (Fig. 2e Extended Data Fig. 2) consistent with previous reports^14^ and our proteomic data (Fig. 1). Although NETosis has been reported in human pulmonary TB tissues^14^, the spatial distribution of NETosis markers across the metabolic landscape of necrotic lesions is unknown. In this regard, we found the majority of GLUT1 expression to be immediately adjacent to the necrotic core in the area comprised of regions 2 and 3 shown in Fig. 2a (Fig. 2f; Extended Data Fig. 2), consistent with the expression of GLUT1 in core-adjacent macrophages within necrotic granulomas in human and NHP TB lungs^43^. Interestingly, this same region contains very few CD66b^+^ neutrophils and is essentially devoid of NETosis markers MPO, NE, and H3-Cit. In contrast, GLUT3 immunostaining was distal to the necrotic core, located within region 4 in Fig. 2a. and extending outward into the adjacent tissue (Fig. 2g; Extended Data Fig. 2). This spatially distinct expression of GLUT1 and GLUT3 (Fig. 2h) suggests region-specific regulation of glucose uptake within the TB granuloma microenvironment. Strikingly, GLUT3 colocalizes with NETosis markers MPO and H3-Cit (Fig. 2i) and MPO, H3-Cit, and NE (Extended Data Fig. 2). Compared to GLUT1 and other Class I glucose transporters, GLUT3 has the highest affinity for glucose^44,45^. Although GLUT3 expression has been observed in necrotic and hypoxic zones of TB granulomas in NHPs^43^, its colocalization with NETosis markers suggests that GLUT3-mediated glucose metabolism may play a role in promoting NETosis and tissue necrosis in human pulmonary TB.

Human necrotic granulomas express GLUT1 directly adjacent to the necrotic core (Fig. 2f, Extended Data Fig. 2,3) in a region that contains CD68+ macrophages (Extended Data Fig. 3). This area may parallel the "hypoxic zone" described in non-human primates, which is composed primarily of GLUT1-expressing macrophages^46^. Within human necrotic TB granulomas, we observed a consistent spatial separation of GLUT1 and GLUT3 expression (Fig. 2f,g, Extended Data Fig. 2, Extended Data Fig. 3d-f). While CD68+ macrophages in the peri-necrotic region colocalize with GLUT1 (Extended Data Fig. 3g,j), there is minimal colocalization with GLUT3 in this region (Extended Data Fig. 3h,k). Notably, a narrow zone of GLUT1- and GLUT3-negative macrophages is present between the core-adjacent GLUT1-positive macrophages and the peripheral GLUT3-positive region (Extended Data Fig. 3i,l). The non-overlapping expression of GLUT1 and GLUT3 suggests the presence of a metabolic gradient across the granuloma that regulates nutrient availability and/or utilization that affects the immune response. In addition to its expression in core-adjacent macrophages, GLUT1 is likely expressed by other cell types within this niche (Extended Data Fig. 3j).

In human non-necrotic granulomas, H&E and DAPI staining confirmed that these lesions contain little, if any, central necrosis (Extended Data Fig. 4a,b). These lesions were composed primarily of CD68+ macrophages (Extended Data Fig. 4c) with CD66b+ neutrophils dispersed throughout the granuloma (Extended Data Fig. 4d).

In contrast to necrotic granulomas, we observed little spatial segregation of GLUT1 and GLUT3 expression (Extended Data Fig. 4e-h) with considerable colocalization in CD66b+ neutrophils (Extended Data Fig. 4i,j) and CD68+ macrophages (Extended Data Fig. 4d-f,k,l) indicating the absence of metabolically segregated zones.

In conclusion, our findings demonstrate that NETosis is a prominent, spatially organized feature of human necrotic TB granulomas, with NETosis markers and GLUT3 strongly colocalizing at the granuloma–tissue interface (Fig. 2, Extended Data Fig. 4k, l). In necrotic granulomas, GLUT-negative macrophages occupy a distinct zone between peri-necrotic GLUT1-positive regions and the distal GLUT3 expression associated with NETotic neutrophils, indicating a structured metabolic organization within the granuloma (Extended Data Fig. 3). In contrast, in non-necrotic granulomas, GLUT1 and GLUT3 expression was colocalized across infiltrating immune cell types, suggesting a link between metabolic compartmentalization and lesion progression. The localization of NETosis markers is consistent with our proteomic data (Fig. 1) and strongly suggests a role for neutrophils in the establishment or maintenance of caseous granulomas. Given its high affinity for glucose, GLUT3 may represent a critical metabolic checkpoint, enabling NETotic neutrophils to sustain glucose uptake under hypoxic or glucose-restricted conditions^43,47^. Together, these findings provide new insight into the metabolic and spatial regulation of NETosis in TB granulomas and highlight GLUT3-mediated glucose metabolism as a potential therapeutic target to modulate TB immunopathology.

### Glucose enhances the viability and effector responses of *Mtb*-infected human neutrophils

The availability and utilization of carbon sources and other nutrients influence the immune responses of macrophages and CD8⁺ T-cells^48,49^. However, whether neutrophil immunometabolism is reprogrammed during *Mtb* infection remains unclear. To determine whether neutrophil viability and response to *Mtb* infection *in vitro* are dependent on nutrient type, neutrophils isolated from healthy donors were cultured with carbon sources known to modulate immune responses, including those of neutrophils^32,50^ (Fig. 3a). Maximum neutrophil viability was observed in medium containing 10 mM glucose compared to the same concentration of other carbon sources at 3, 6, 9, 12, and 15 hours after plating (Fig. 3b). *Mtb* infection reduced neutrophil viability overall (Extended Data Fig. 5); however, 10 mM glucose supported maximum viability at 3, 6, and 9 hours post-infection (Fig. 3c). We next assessed whether *Mtb* infection also alters neutrophil effector functions, including MPO activity, NE activity, and NETosis, and whether these responses are dependent on nutrient type. Following *Mtb* infection, NE and MPO activities in the culture medium (Fig. 3d,e) and the number of NETosis-positive cells (Fig. 3f) were significantly higher when neutrophils were cultured in 10 mM glucose compared to other carbon sources at the same concentration. These findings, along with our proteomic (Fig. 1) and mIF data (Fig. 2), suggest that the availability and utilization of glucose within the spectrum of human TB lesions may be key determinants of neutrophil survival, effector functions, and TB immunopathology.

**Fig. 3.**
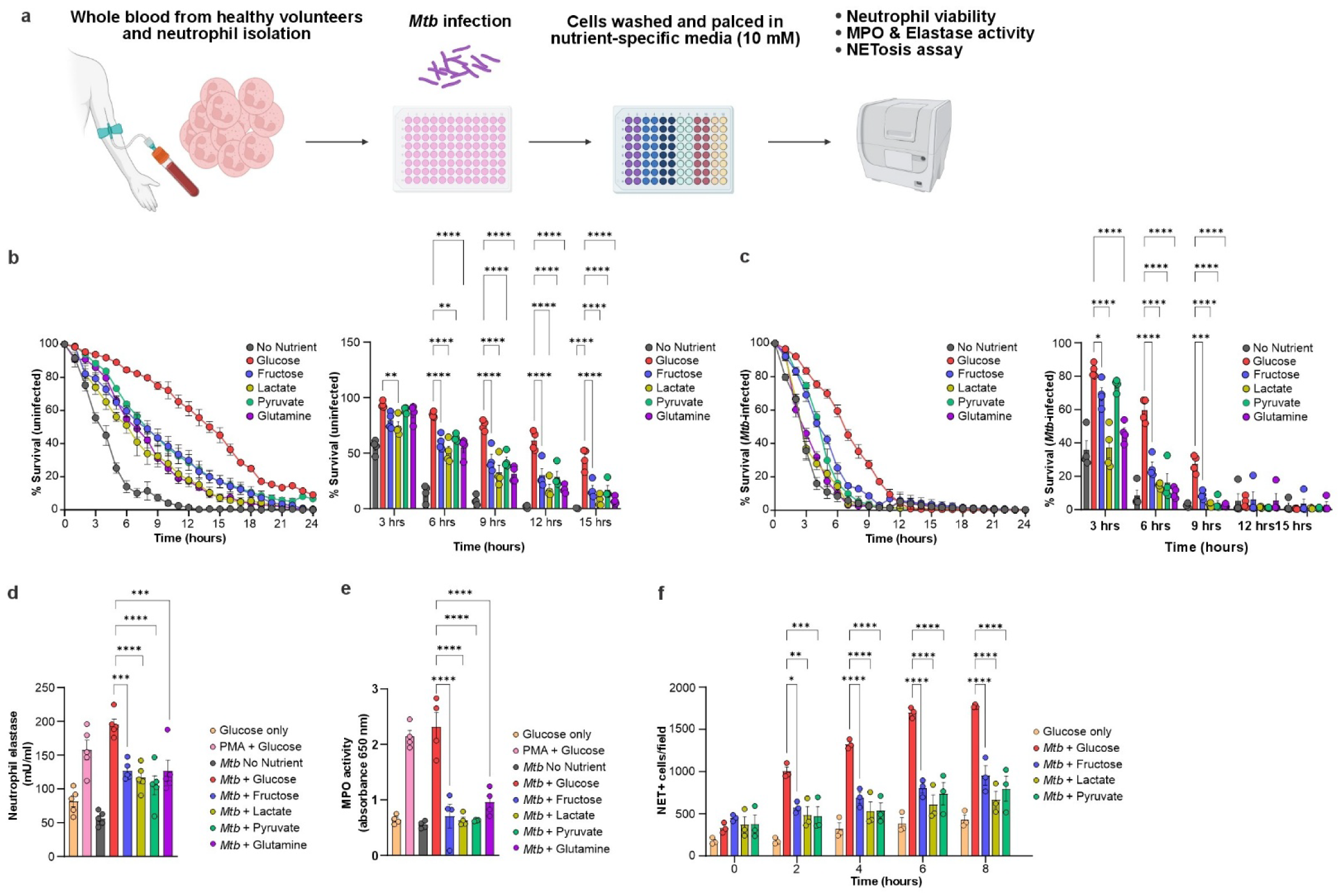
| **Glucose increases viability and effector responses of *Mtb*-infected human neutrophils. a**, Workflow for isolating neutrophils from healthy donors and measuring cell viability and effector functions. **b**, **c**, Survival of uninfected (**b**) and *Mtb*-infected (MOI of 5) (**c**) human neutrophils cultured with no carbon source or with 10 mM glucose, fructose, lactate, pyruvate, or glutamine. Column graphs in (**b**) and (**c**) show neutrophil viability at 3, 6, 9, 12, and 15 hours with and without infection. Data are shown as mean ± SEM; n = 4 technical replicates per group. Statistical significance was determined using two-way ANOVA with Tukey’s multiple comparisons test (\**P* < 0.03, \*\**P* < 0.002, \*\*\**P* < 0.0002, \*\*\*\**P* < 0.0001). **d**, Neutrophil elastase activity in neutrophil culture medium containing various nutrients (10 mM) and infected with *Mtb* (MOI of 5). Controls include uninfected and PMA-treated (100 ng/mL) neutrophils cultured in 10 mM glucose. Data are shown as mean ± SEM; n = 5 technical replicates per group. Statistical analysis was performed using one-way ANOVA with Dunnett’s multiple comparisons test (\*\*\**P* < 0.0002, \*\*\*\**P* < 0.0001). **e**, MPO activity in neutrophil culture medium containing various nutrients (10 mM) and infected with *Mtb* (MOI of 5). Controls include uninfected and PMA-treated (100 ng/mL) neutrophils cultured in 10 mM glucose. Data are shown as mean ± SEM; n = 5 technical replicates per group. Statistical analysis was performed using one-way ANOVA with Dunnett’s multiple comparisons test (\*\*\*\**P* < 0.0001). **f**, NETosis in neutrophils cultured with various nutrients (10 mM) and infected with *Mtb* (MOI 1:5). Columns are representative of 3 independent experiments. NET-positive cells were identified by extracellular DNA release and quantified using Gen5 software on the Agilent BioTek Cytation-5 cell imaging multimode reader. All assays were performed independently at least 3 times.

### *Mtb* triggers an immediate, pathogen-specific, glucose-dependent oxidative burst in human neutrophils

Neutrophils can respond to microbial threats by the rapid production and release of superoxide anions (O ^●─^), in a process known as the oxidative burst. The conversion of O_2_ to O ^●─^ by NOX2 is fueled by NADPH, which is generated through the PPP^31,51^. Traditionally, ROS species including hydrogen peroxide (H_2_O_2_), the hydroxyl radical (OH^●^), hypochlorous acid (HOCl), etc., have been used as proxies for measuring the pathogen-induced neutrophil oxidative burst. However, the ability to accurately measure these diverse ROS species is significantly affected by probe specificity, pH, oxygen level, and cell permeability^52^. Further, conventional ROS assays cannot distinguish between ROS generated by the electron transport chain (ETC) and those produced via NOX2 activity during the oxidative burst, complicating data interpretation^52^. Therefore, we quantified neutrophil oxidative burst responses directly by measuring the oxygen consumption rate (OCR) of NOX2 in real time using the Agilent Seahorse XFe96 Extracellular Flux (XF) Analyzer^29,31^ within BSL-3 containment. By pharmacologically inhibiting oxygen consumption attributed to oxidative phosphorylation (OXPHOS), we were able to accurately quantify the oxygen consumption resulting solely from the neutrophil oxidative burst. To the best of our knowledge, use of the XF Analyzer to measure neutrophil oxidative burst in response to live or pathogenic bacteria has not been reported.

Although we found no difference in glucose content between granulomatous and non-TB lung tissues (Extended Data Fig. 1a,b), NETosis markers and GLUT-3 levels were elevated in necrotic regions (Fig.1), suggesting that nutrient utilization within the TB granuloma microenvironment may influence neutrophil effector responses. To determine whether the fuel source can affect the oxidative burst, we used the XF Analyzer to measure the oxidative burst in neutrophils isolated from healthy donors (Fig. 4a). As described previously^29^, rotenone and antimycin A were injected first to inhibit oxygen consumption by OXPHOS, *i.e*., the terminal oxidase complexes I and II of the ETC. A stimulant, such as PMA, is then injected into the medium through the second port and total NOX2-mediated oxygen consumption is quantified by calculating the area under the OCR curve (AUC) (Fig. 4b). We further assessed neutrophil bioenergetics by measuring the extracellular acidification rate (ECAR), a proxy indicator of glycolysis, and oxygen consumption rate (OCR), an indicator of mitochondrial (OXPHOS) metabolism, following the same injection strategy applied to macrophages^26,53^ (Fig. 4b).

**Fig. 4.**
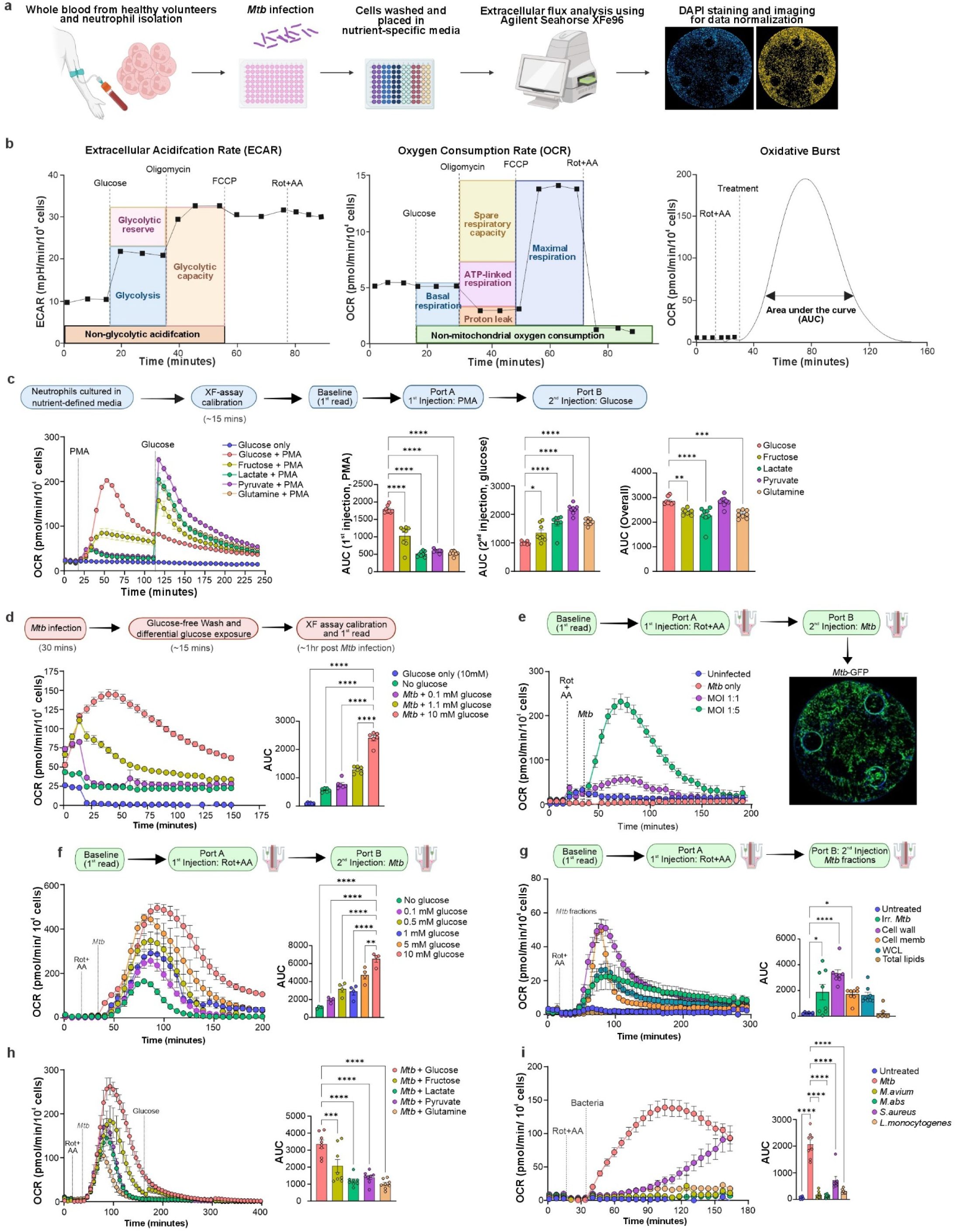
| **Intact *Mtb* triggers an immediate, glucose-dependent oxidative burst in human neutrophils**. **a**, Workflow for isolating neutrophils from healthy donors and measuring neutrophil oxidative burst and other bioenergetic parameters during *Mtb* infection using the Agilent Seahorse XFe96 Extracellular Flux Analyzer. All XF data were normalized to 10,000 cells using Gen5 software on an Agilent BioTek Cytation 5 Multimode Reader. Serum-free RPMI 1640 medium was used in all assays. **b**, Representations of XF assay design, including injection strategies for measuring extracellular acidification rate (ECAR), oxygen consumption rate (OCR), and oxidative burst. The magnitude of oxidative burst is determined by calculating the area under the OCR curve (AUC) and is plotted as column graphs. **c**, Oxidative burst of neutrophils initially cultured on glucose (10 mM; Glu and Glu + PMA groups) or other nutrients (10 mM) following stimulation with PMA. After calibration and three baseline measurements, PMA (100 ng/mL final concentration) was injected via the first injection port and OCR was measured. Approximately 100 min later, glucose (10 mM final concentration) was injected via the second port (final glucose concentration for Glu and Glu + PMA groups was 20 mM, all others were 10 mM) and OCR measured. The magnitude of the initial oxidative burst (UAC 1^st^ injection, PMA) was determined from the area between the first and second injections. The second oxidative burst (AUC 2nd injection, glucose) was determined from the AUC following the second injection. The overall oxidative burst (AUC Overall) is the sum of the two bursts. **d**, Neutrophil oxidative burst following *Mtb* infection (MOI of 5). Neutrophils cultured in medium containing 10 mM glucose were exposed to *Mtb* for 30 minutes, washed three times in glucose-free medium to remove extracellular bacilli, then cultured in medium containing 0-10 mM glucose. Uninfected controls were maintained in 10 mM glucose. AUC was calculated beginning with the first OCR measurement. **e**, Infection of neutrophils by injecting *Mtb* bacilli through XF injection port B in medium containing 10 mM glucose. After calibration and baseline measurements, rotenone (1.25 μM) and antimycin A (2.5 μM) were injected via port A to inhibit mitochondrial respiration. *Mtb*-GFP bacilli were injected through port B at an MOI of 1 or 5. After the completion of the XF run, *Mtb*-GFP distribution in the wells was visualized using the Agilent BioTek Cytation 5 Cell Imaging Multimode Reader to confirm uniform bacterial distribution. **f**, Oxidative burst of neutrophils cultured on glucose (0-10 mM) following injection of *Mtb* bacilli (MOI of 5) via port B. **g,** Oxidative burst of neutrophils cultured on glucose (10 mM) in response to live *Mtb* (MOI of 5), irradiated (irr.) *Mtb*, *Mtb* whole-cell lysate, or *Mtb* cellular fractions containing cell wall, cell membrane, or total lipids. **h**, Oxidative burst of neutrophils in medium containing various nutrients, with rotenone (1.25 μM) and antimycin A (2.5 μM) injected via port A prior to *Mtb* injection (MOI 1:5) via Port B. Glucose (10 mM final concentration) was later injected into all wells. **i**, Oxidative burst of neutrophils cultured in 10 mM glucose following infection with *Mtb*, *M. avium*, *M. abscessus*, *S. aureus*, or *L. monocytogenes* (all MOI of 5), via injection from port B. **c–i,** column graphs represent the magnitude of the oxidative burst and are shown as the mean ± SEM of the areas under the OCR curve for each group (n = 6-8 technical replicates per group). Statistical significance was determined using ordinary two-way ANOVA with Tukey’s multiple comparisons test (\**P* < 0.03, \*\**P* < 0.002, \*\*\**P* < 0.0002, \*\*\*\**P* < 0.0001). All assays were performed independently three or more times.

First, neutrophils were cultured in media containing various carbon sources at 10 mM, PMA was injected into the culture media, and the magnitude of the oxidative burst was quantified by calculating the area under the OCR curve from the time of PMA injection to the time of the second injection (Fig. 4c). Neutrophils cultured in 10 mM glucose exhibited the highest oxidative burst response followed by fructose, with cells grown on other carbon sources exhibiting small, but significant, oxidative responses (Fig. 4c, AUC 1^st^ injection, PMA). Importantly, previous studies, including those examining metabolic partitioning of the burst, describe only the temporal phases of a single activation and do not address whether multiple, discrete bursts can occur^54^. Thus, whether the neutrophil oxidative burst can be dynamically reactivated is an unresolved question. To investigate this, we measured the OCR of PMA-exposed neutrophils from the time of glucose injection until the end of the assay and calculated the oxidative burst response as the AUC (Fig. 4c OCR, AUC 2^nd^ injection, glucose). Notably, neutrophils initially cultured with non-glucose carbon sources exhibited a pronounced second oxidative burst in response to the addition of glucose, suggesting a metabolically regulated, hierarchical activation. This provides strong evidence that glucose is a critical metabolic fuel for maximal neutrophil oxidative responses, and that addition of glucose can metabolically retrigger glucose-naïve PMA-activated neutrophils (Fig. 4c, AUC 2^nd^ injection, glucose). In contrast, neutrophils initially cultured on 10 mM glucose did not exhibit a second burst, indicating that a glucose-fueled, PMA-induced oxidative burst is a rapid, one-time event (Fig. 4c, AUC 2^nd^ injection, glucose; AUC [Overall]).

Neutrophils have been shown to rapidly phagocytose, but not eliminate, *Mtb*^55^. To determine whether *Mtb*-infection of human neutrophils rapidly induces an oxidative burst, we measured the OCR of neutrophils following *Mtb* infection. Neutrophils were cultured with various nutrients and infected with *Mtb* at an MOI of 5, and oxygen consumption by OXPHOS inhibited by rotenone and antimycin A. After exposure to *Mtb* for 30 minutes, neutrophils were washed to remove unbound bacilli and fresh media containing glucose at various concentrations was added. Intriguingly, XF analysis revealed that an *Mtb*-induced, glucose concentration-dependent oxidative burst had begun prior to the first OCR measurement (Fig. 4d; note y-axis), suggesting rapid activation of NOX2.

To quantify the entire *Mtb*-induced oxidative burst, we modified our XF assays so that live *Mtb* bacilli are directly injected into the culture medium through the second XF port following the co-injection of rotenone and antimycin-A. Strikingly, *Mtb* injection rapidly triggered an oxidative burst, indicating an immediate, contact-dependent neutrophil response (Fig. 4e). Importantly, we confirmed that injection of bacilli provides uniform bacterial distribution across XF culture wells by imaging *Mtb*-GFP following XF analysis (Fig. 4e). Using direct *Mtb* injection, we repeated the glucose concentration experiment (Fig. 4d) and observed a virtually immediate concentration-dependent oxidative burst with 10 mM glucose supporting the most robust response (Fig. 4f) as seen previously (Fig. 4d). Next, we next sought to determine whether the *Mtb*-induced oxidative burst is dependent on *Mtb* viability or specific structural components of the bacillus. Using the same injection approach, neutrophils were exposed to live or irradiated *Mtb, Mtb* whole cell lysate, or bacillary fractions containing *Mtb* cell membrane, cell wall, or lipids (Fig. 4g, Extended Data Fig. 6). Consistent with Fig. 4e, live *Mtb* elicited the most pronounced oxidative burst compared to irradiated *Mtb* or *Mtb* fractions (Extended Data Fig. 6). Interestingly, exposure of neutrophils to irradiated *Mtb* or fractions containing cell wall or membrane components resulted in a statistically significant oxidative burst (Fig. 4g). *Mtb* whole cell lysate induced an oxidative response that did not reach statistical significance, while exposure of neutrophils to *Mtb* lipids did not generate a measurable response (Fig. 4g).

We next asked whether a maximal *Mtb*-induced oxidative burst occurs in neutrophils cultured on glucose, as was observed with PMA (Fig. 4c). Consistent with our results using PMA,10 mM glucose supported the most robust *Mtb*-induced oxidative burst, followed by fructose, lactate, pyruvate, and glutamine at the same concentration (Fig. 4h). However, in contrast to the PMA-induced oxidative response in the presence of various carbon sources (Fig. 4c), injection of glucose following the *Mtb*-induced oxidative response did not elicit a second, increased burst in neutrophils cultured on other carbon sources. This indicates that the *Mtb*-induced oxidative burst follows a mechanism distinct from that of non-physiological PMA (Fig. 4h). Unlike PMA-stimulated neutrophils, *Mtb*-infected neutrophils become metabolically unresponsive to glucose following the initial oxidative burst and cannot be metabolically retriggered by subsequent glucose addition.

Finally, to ascertain whether the oxidative burst induced by *Mtb* is a general response to bacterial infection or specific to *Mtb*, we infected neutrophils with non-tuberculous mycobacteria (NTM) *Mycobacterium avium* (*Mav*) or *Mycobacterium abscessus* (*Mab*), or intracellular bacteria *Staphylococcus aureus* (*Sa*) or *Listeria monocytogenes* (*Lm*). Remarkably, only *Mtb* rapidly elicited a robust oxidative burst, whereas *Mav, Mab* and *Lm* induced, little, if any, response. Notably, *Sa* elicited a slower, gradual oxidative response compared to that induced by *Mtb* (Fig. 4i).

In sum, our results demonstrate that *Mtb* bacilli can rapidly trigger a contact-dependent oxidative burst in human neutrophils and that glucose is required to produce a maximal response. *Mtb*-exposed neutrophils cultured on non-glucose carbon sources produce a single oxidative burst and are unresponsive to subsequent addition of glucose, indicating that *Mtb*-driven oxidative responses involve a metabolic program that is distinct from PMA-induced activation. Of the pathogenic bacteria tested, only *Mtb* elicits an immediate and robust oxidative burst. This finding highlights the distinctive ability of *Mtb* to activate human neutrophils and reveals a previously unrecognized aspect of neutrophil physiology. The use of XF analysis with direct *Mtb* injection enabled precise, real-time measurement of this response, revealing both the temporal resolution and metabolic requirements of *Mtb*-induced neutrophil activation, which represents a substantial technical advancement.

### *Mtb* rapidly triggers an oxidative burst in human neutrophils through Dectin-1 and Fcγ receptors

The virtually immediate oxidative burst following exposure to live *Mtb* (Fig. 4) strongly suggests a contact-dependent process involving cell surface pattern recognition receptors (PRRs). Neutrophils isolated from patients with pulmonary TB exhibit increased expression of Toll-like receptors (TLR2 and TLR4) and Fc gamma receptors (FcγRs) have been shown to influence immune activation during *Mtb* infection^56^. Interestingly, increased PRR expression is linked to reduced phagocytic capacity, suggesting the possibility that these receptors regulate neutrophil functions beyond phagocytosis, including oxidative burst activity^56^. In addition, C-type lectin receptors (CLRs) such as macrophage inducible Ca²⁺-dependent lectin receptor (Mincle, CLEC4E) and Dectin-1 (CLEC7A) recognize *Mtb* ligands and trigger downstream signaling through spleen tyrosine kinase (Syk) and mitogen-activated protein kinase (MAPK) pathways that drive ROS production^57,58^. However, whether these PRRs contribute to *Mtb*-induced oxidative burst in human neutrophils is unknown.

To identify PRRs involved in the *Mtb*-induced oxidative burst, neutrophils were incubated with TLR2 or TLR4 inhibitors or blocking antibodies directed against Mincle, Dectin-1, FcγRI/CD64, FcγRII/CD32, or FcγRIII/CD16 prior to injection of *Mtb* and OCR measurements (Fig. 5a). Exposure of neutrophils to the TLR2 inhibitor TLR2-IN-C29^59^ (Fig. 5b,c; Extended Data Fig. 7a) or the TLR4 inhibitor TAK-242^60^ (Fig. 5d,e; Extended Data Fig. 7b) did not significantly alter the oxidative burst in response to *Mtb*. Similarly, incubation of neutrophils with a blocking antibody directed against Mincle^61^ did not affect the oxidative burst response to *Mtb* (Fig. 5d, e; Extended Data Fig. 7c). In contrast, addition of a Dectin-1 blocking antibody^62^ resulted in a marked, dose-dependent reduction in the *Mtb*-induced oxidative burst (Fig. 5d,e; Extended Data Fig. 7d), consistent with the identification of *Mtb* α-glucan polysaccharides as a Dectin-1 ligands^63^. Similarly, addition of blocking antibodies against CD16^64^ (Fig. 5f,g; Extended Data Fig. 7e), CD32^65^ (Fig. 5f,g; Extended Data Fig. 7f), or CD64^66^ (Fig. 5f, g; Extended Data Fig. 7g) significantly reduced the *Mtb*-induced oxidative burst in a dose-dependent manner (Extended Data Fig. 7h). It is important to note that Fcγ receptor interactions are classically linked to antibody-dependent mechanisms and can shape neutrophil antimicrobial responses during *Mtb* infection^67^. However, the oxidative burst described here was observed under serum-free conditions, arguing against a requirement for IgG opsonization.

**Fig. 5.**
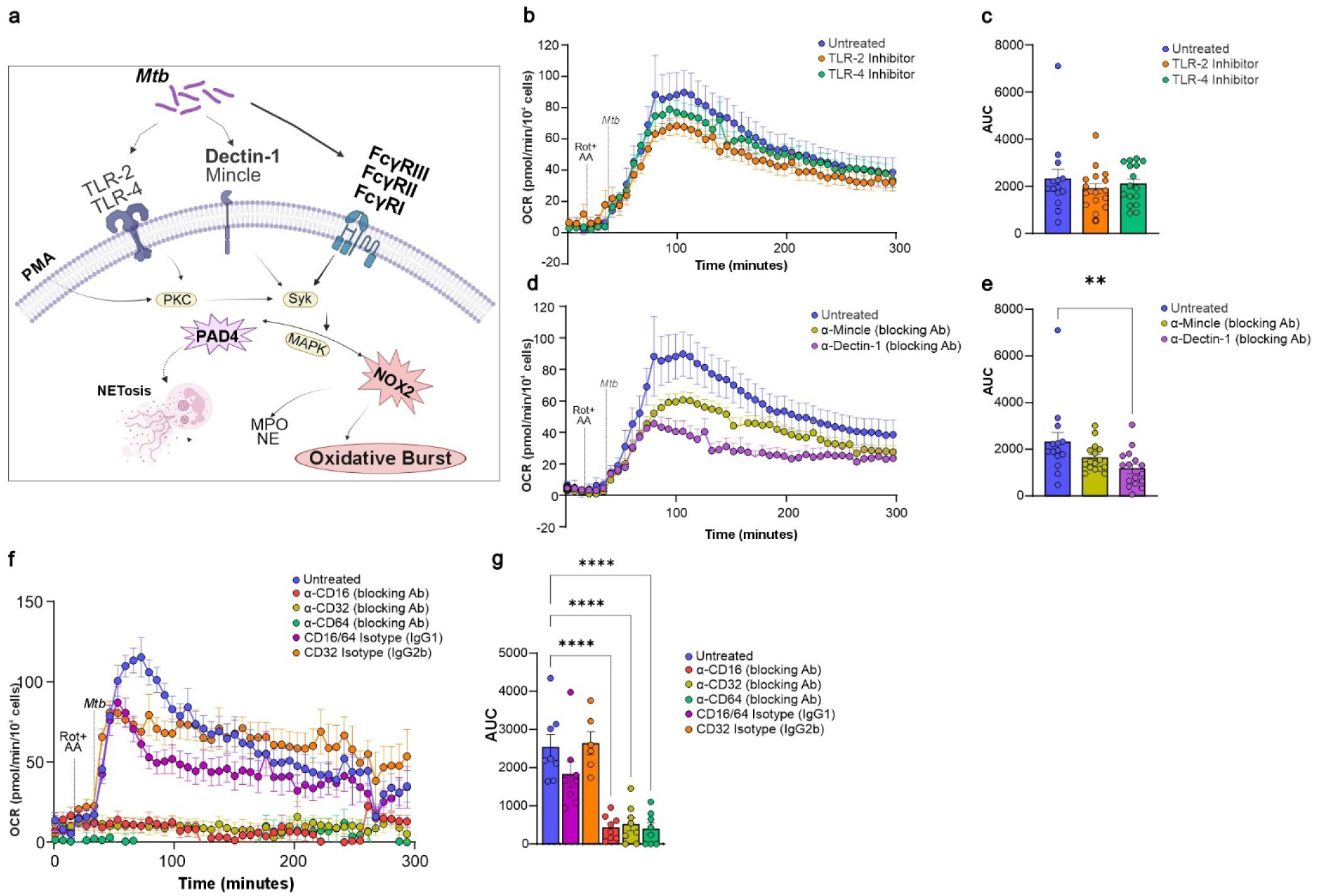
| ***Mtb* rapidly triggers an oxidative burst in human neutrophils through Dectin-1 and Fcγ receptors. a**, Representation of *Mtb*-induced neutrophil activation via surface pathogen recognition receptors (PRRs), including Toll-like receptors (TLRs), C-type lectin receptors (CLRs), and Fc gamma receptors (FcγRs). *Mtb* triggers oxidative burst through receptor-mediated signaling pathways leading to the generation of reactive oxygen species (ROS), activation of MPO and NE, and NETosis. In contrast, PMA bypasses PRRs and directly activates protein kinase C (PKC) to stimulate NOX2. **b**-**g**, Neutrophils were obtained from healthy donors and the oxidative burst was measured using the Agilent Seahorse XFe96 Extracellular Flux Analyzer. All XF data were normalized to 10,000 cells using Gen5 software on an Agilent BioTek Cytation 5 Multimode Reader. Serum-free RPMI 1640 medium was used in all assays. One hour prior the start of the assay, neutrophils were exposed to the TLR2 inhibitor TLR2-IN-C29 (100ng/mL) or TLR4 inhibitor TAK 22 (100 ng/mL) (**b, c**), or blocking antibodies directed against Mincle (1 μg/mL) or Dectin-1 (1 μg/mL) (**d**,**e**), or CD16, CD32, or CD64 and isotype control antibodies at 1 μg/mL (**f**, **g**). Rotenone (1.25 μM) and antimycin A (2.5 μM) were then injected via port A to inhibit mitochondrial respiration, followed by *Mtb* injection via port B (MOI of 5). Column graphs represent the magnitude of the oxidative burst and are shown as the mean ± SEM of the areas under the OCR curve for each group (n = 6-8 technical replicates per group). Statistical significance was determined using ordinary one-way ANOVA with Dunnett’s multiple comparisons test (\**P* < 0.03, \*\**P* < 0.002, \*\*\**P* < 0.0002, \*\*\*\**P* < 0.0001). All assays were performed independently three or more times.

Together, these data demonstrate that *Mtb* induces a robust neutrophil oxidative burst through a mechanism that involves Dectin-1 and Fcγ receptors, even in the absence of serum, suggesting antibody-independent signaling^68^. Also, the neutrophil oxidative burst relies on integrated signals from multiple PRRs. Lastly, given that Dectin-1-deficient mice exhibit markedly reduced neutrophil recruitment to *Mtb*-infected lungs^63^, our findings suggest that Dectin-1 engagement is critical for initiating the metabolic reprogramming and oxidative burst in neutrophils that contribute to tissue damage.

### *Mtb* infection triggers glucose-driven bioenergetic reprogramming in neutrophils and a GM-CSF-enhanced oxidative burst

Neutrophils are the immune system’s first responders to bacterial pathogens. However, the magnitude of neutrophil responses at the site of infection is influenced by host-derived cytokine and chemokine signals, as well as pathogen-associated cues, including the *Mtb* cell wall glycolipid lipoarabinomannan (LAM), in a phenomenon known as neutrophil priming^69^. During *Mtb* infection, GM-CSF, TNF-α, and IFN-γ can prime neutrophils, thereby enhancing their antimicrobial functions to promote bacterial clearance^70,71^. However, if primed neutrophil responses persist, as often occurs in chronic TB, they can drive excessive oxidative burst, increased ROS production, and NETosis, contributing to immunopathology (Fig. 6a)^72,73^. To investigate the effects of cytokine priming on the oxidative burst response to *Mtb*, human neutrophils were cultured in medium containing GM-CSF, TNF-α, or IFN-γ, exposed to *Mtb*, and the oxidative burst was measured. Addition of IFN-γ had no effect on the *Mtb*-induced oxidative burst (Fig. 6b). Interestingly, priming neutrophils with TNF-α significantly reduced the oxidative burst (Fig. 6c), whereas GM-CSF priming markedly enhanced this response (Fig. 6d), illustrating cytokine-specific modulation of neutrophil oxidative activity.

**Fig. 6.**
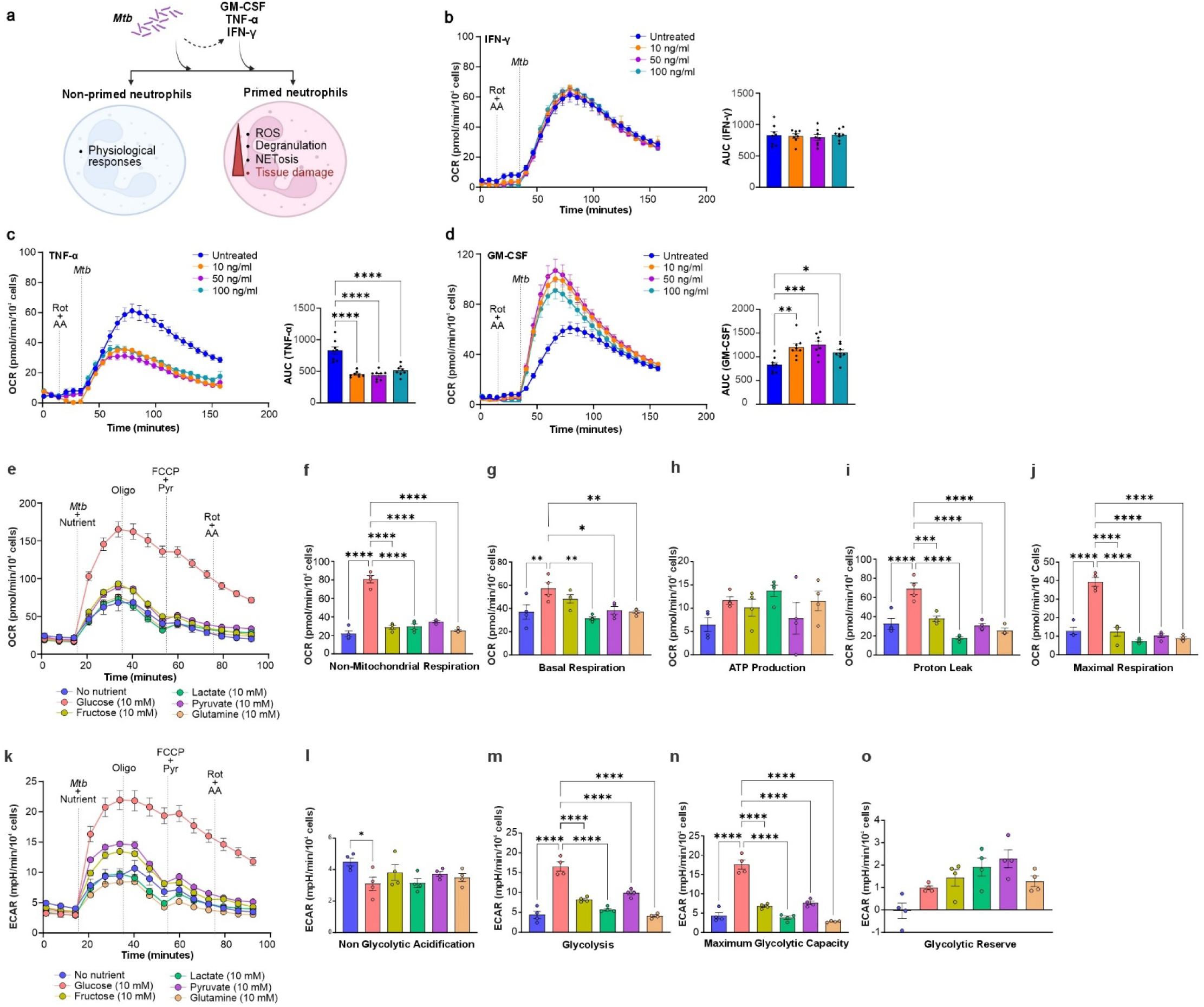
| ***Mtb* infection triggers glucose-driven bioenergetic reprogramming in neutrophils and a GM-CSF-enhanced oxidative burst. a,** Illustration of *Mtb*-induced neutrophil priming and effector functions. Under homeostatic conditions, unprimed neutrophils carry out physiological immune surveillance with limited activation. During *Mtb* infection, elevated levels of inflammatory cytokines such as GM-CSF, TNF-α, and IFN-γ contribute to neutrophil priming. Upon encountering pathogens at the site of infection, primed neutrophils exhibit amplified effector functions including production of reactive oxygen species (ROS), degranulation, and NETosis. While these responses are critical for pathogen clearance, their dysregulation can result in excessive tissue damage and contribute to tissue damage. **b**–**d**, *Mtb*-induced oxidative burst in neutrophils exposed to inhibitors of IFN-γ (**b**) TNF-α (**c**) or GM-CSF (**d**) at 10, 50 and 100 ng/mL for 1 hour prior to infection. Oxidative burst was measured using the Agilent Seahorse XFe96 Extracellular Flux Analyzer. Rotenone (1.25 μM) and antimycin A (2.5 μM) were injected via port A to inhibit mitochondrial respiration, followed by *Mtb* injection via port B (MOI of 5). All XF data were normalized to 10,000 cells using Gen5 software on the Agilent BioTek Cytation 5 Cell Imaging Multimode Reader. Column graphs represent the magnitude of the oxidative burst and are shown as the mean of the areas under the OCR curve for each group. **e**–**j**, The OCR (**e**) in *Mtb*-infected human neutrophils (MOI of 5) cultured with various nutrients (10 mM), and other parameters of mitochondrial function including non-mitochondrial respiration (**f**), basal respiration (**g**), ATP production (**h**), proton leak (**i**), and maximal respiration (**j**) were determined as outlined in Fig. 4b. **k-i,** The extracellular acidification rate (ECAR) **(k**) in *Mtb*-infected human neutrophils (MOI of 5) cultured with various nutrients (10 mM), and non-glycolytic acidification (**I**), glycolysis (**m**), maximum glycolytic capacity (**n**), glycolytic reserve (**o**) was determined. Dashed lines indicate injections starting with *Mtb* (MOI of 5) plus nutrients, followed by oligomycin (1.5 μM), FCCP (1.5 μM) plus pyruvate (1mM) and lastly rotenone (1.25 μM) plus antimycin A (2.5 μM). ECAR and OCR parameters were calculated from the same assay as previously described by Bossche, *et. al*^53^. Data are presented as mean ± SEM (n = 6-8 technical replicates per group). Statistical significance was determined using ordinary one-way ANOVA with Dunnett’s multiple comparisons test (\**P* < 0.03, \*\**P* < 0.002, \*\*\**P* < 0.0002, \*\*\*\**P* < 0.0001). All assays were performed independently three or more times.

We next examined whether neutrophil energy production is influenced by nutrient availability and/or exposure to *Mtb*. We measured the extracellular acidification rate (ECAR) as a proxy for glycolysis, and the OCR to monitor mitochondrial respiration (OXPHOS) (Fig. 4b) using a previously reported XF analysis strategy^26,53^, with the exception that here, *Mtb* and carbon sources were injected first. Consistent with previous reports^74^, exposing neutrophils to *Mtb* caused a rapid increase in mitochondrial respiration (measured as OCR, Fig. 6e-j) and glycolysis (measured as ECAR, Fig. 6k-o) in a carbon-source dependent manner. Like the oxidative burst response (Fig. 4h), neutrophil bioenergetic parameters were significantly elevated in the presence of glucose compared to other nutrients (Fig. 6e-h,j,k,m,n). Using a modified XF assay that integrates mitochondrial and glycolytic readouts^53^, non-mitochondrial respiration (Fig. 6f), basal respiration (Fig. 6g), proton leak (Fig. 6i), maximal respiration (Fig. 6j), glycolysis (Fig. 6m), and maximum glycolytic capacity (Fig. 6n) were significantly increased upon *Mtb* infection in glucose-containing medium. In contrast, no differences were observed in ATP production (Fig. 6h), non-glycolytic acidification (Fig. 6l), or glycolytic reserve (Fig. 6o) between neutrophils cultured on glucose and other carbon sources.

Since metabolic changes are closely linked to immune responses, we next investigated whether neutrophil cytokine release is nutrient dependent as observed for the oxidative burst, OCR, and ECAR. Interestingly, neutrophil cytokine secretion was largely independent of nutrient type, with the levels of most cytokines, including IL-6, IFN-γ, GM-CSF, TNF-α, increasing upon *Mtb* infection regardless of the fuel source (Extended Data Fig. 8).

Overall, these results demonstrate that *Mtb* infection induces distinct metabolic reprogramming in neutrophils, marked by glucose-driven increases in glycolysis and OXPHOS. Also, *Mtb*-infected neutrophils display divergent responses to cytokine priming. For example, GM-CSF significantly augments the oxidative burst, consistent with classical priming of NADPH oxidase. However, this also argues that a GM-CSF-rich microenvironment, such as the *Mtb*-infected lung, could potentiate the damaging, glucose-fueled neutrophil oxidative burst. In contrast, priming neutrophils with TNFα reduced the oxidative burst, whereas priming with IFNγ had no detectable effect, indicating cytokine-specific regulation of neutrophil oxidative metabolism. Lastly, while glucose utilization enhances neutrophil bioenergetic responses such as the oxidative burst, OCR, and ECAR, cytokine production is largely fuel-independent, revealing an immunometabolic plasticity that uncouples metabolic signaling from cytokine secretion during *Mtb* infection.

### *Mtb* infection rewires neutrophil metabolic flux from glycolysis towards the PPP for rapid oxidative burst

Neutrophils have high energy requirements to support their effector functions, including the NOX2-mediated oxidative burst, ROS production, and NETosis, which are thought to be heavily dependent on glycolysis^75^. However, during PMA stimulation or physiological stresses like hypoxia, nutrient deprivation, or COPD-related inflammation, neutrophils exhibit metabolic plasticity and can redirect their metabolism to the PPP and gluconeogenesis, and utilize non-glucose substrates like glutamine and glycogen^31,32^. Given their metabolic flexibility under stress, we sought to understand how neutrophils alter their metabolic phenotype upon *Mtb* infection. As a first step, we performed RNA sequencing (RNA-seq) analysis to capture transcriptional changes that occur within one hour following *Mtb* infection. Principal component analysis (PCA) of transcriptomic data shows distinct clustering between *Mtb*-infected and uninfected neutrophils, indicating infection-associated transcriptional reprogramming (Extended Data Fig. 9f). However, the expression of genes encoding members of key metabolic pathways, including glycolysis, PPP, TCA, OXPHOS, and NETosis was not significantly altered by *Mtb* infection (Extended Data Fig. 9a-e). The minimal transcriptional response we observed is consistent with prior work showing that activated human neutrophils can rapidly change phenotype through signal-dependent translation of constitutively expressed mRNAs, rather than broad de novo transcription^76^. Moreover, this also suggests that core metabolic pathways remain transcriptionally stable, implying that the observed metabolic rewiring occurs because neutrophils are preloaded with all necessary metabolic enzymes.

To identify *Mtb*-induced changes in metabolic flux through glycolysis and the PPP, neutrophils were infected with *Mtb*, washed, and cultured in medium containing ^13^C_1,2_-glucose. PCA analysis of global metabolomic profiles reveals a clear separation between *Mtb*-infected and uninfected neutrophils (Extended Data Fig. 9g) indicating infection-associated metabolic shifts. Several lines of evidence suggest that the continued, and sometimes enhanced use of ^13^C_1,2_-glucose carbons is consistent with reprogramming towards high flux NADPH generation rather than biosynthesis. Firstly, *Mtb*-infected neutrophils show a coordinated contraction of both glycolytic and PPP metabolites pools, which was evident by a marked reduction in the abundance glycolytic intermediates (G6P/F6P, F-1,6-P, G3P, PGA, PEP, and Pyr; Fig. 7a) as well as PPP metabolites, including erythrose-4-phosphate (E4P) and sedoheptulose-7-phosphate (S7P) (Fig. 7b and Extended Data Fig. 9h). Yet, the fractional ^13^C enrichment either increases (F1,6BP, PEP), decreases (G3P) or remain stable (remaining metabolites), suggesting that the smaller pools are supplied by labeled glucose rather than by pre-existing unlabeled carbon sources such as cellular glycogen, glycerol/lipids, amino acids, etc. Secondly, E4P levels are reduced in *Mtb* infected neutrophils, while their isotopologue distributions from labeled glucose remain similar to uninfected controls. Lastly, since metabolite abundance is not an indicator of relative flux, we quantified relative carbon flux through the oxPPP by calculating the isotopologue distribution of lactate, comparing M+2 and M+3 species derived from glycolysis with M+1 lactate, which is generated predominantly via the PPP, as described^77^. These data, together with shrinking upper and lower glycolytic intermediate levels, show that neutrophils undergo a significant metabolic shift from glycolysis toward increased PPP flux upon *Mtb* infection (Fig. 7c). Previous metabolic tracing experiments using ^13^C_1_- and ^13^C_1,2_-glucose in PMA-stimulated neutrophils demonstrated a comparable metabolic shift toward the oxPPP, which was posited as a means to maximize NADPH production required for NOX2-mediated oxidative burst^31^ (Fig. 7d).

**Fig. 7.**
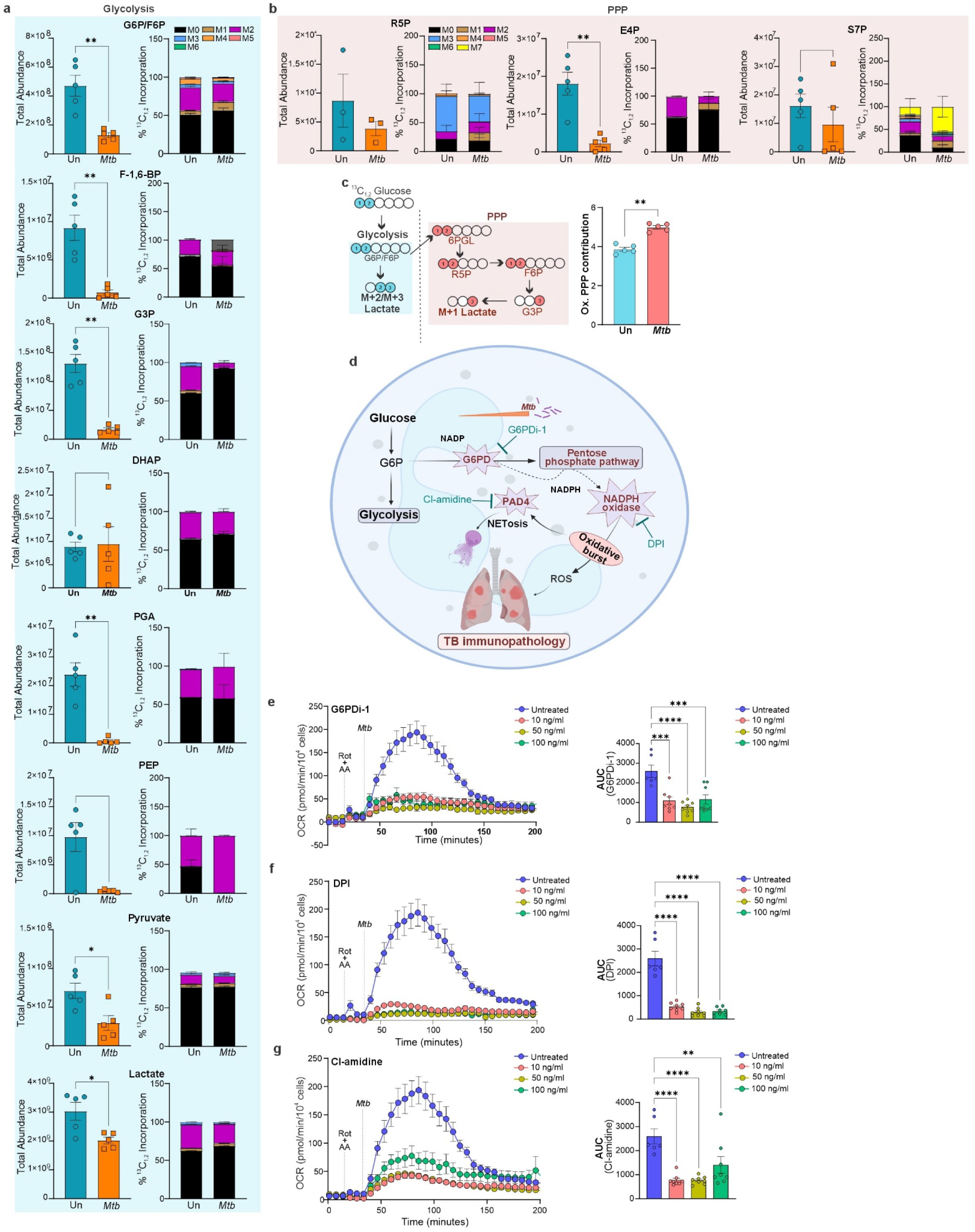
| ***Mtb* infection rewires neutrophil metabolic flux from glycolysis towards the PPP. a**, Total abundance (left) and isotopologue distribution (right) of glycolytic intermediates in uninfected and *Mtb*-infected neutrophils. **b**, Total abundance (left) and isotopologue distribution (right) of PPP intermediates in uninfected and *Mtb*-infected neutrophils. **c**, Schematic representation of lactate isotopologue distribution; glycolysis of ^13^C_1,2_-glucose (blue box) generates M+2 lactate, while PPP activity (pink box) produces M+1 lactate. Column graph (right) shows the oxidative PPP fractional contribution estimated from lactate isotopologue ratios (M+1/M+2) using the formula (M+1/M+2)/(3 + M+1/M+2). **d**, Schematic illustrating the metabolic rewiring of neutrophils upon *Mtb* infection. In steady-state conditions, glucose is metabolized primarily through glycolysis to generate ATP. Upon *Mtb* infection, glucose-6-phosphate (G6P) is diverted into the PPP by the activity of glucose-6-phosphate dehydrogenase (G6PD). The resulting NADPH fuels NOX2-mediated oxidative burst, leading to ROS production, calcium-dependent activation of PAD4, and subsequent NETosis. Together, ROS and NETosis contribute to chronic TB-associated immunopathology. **e**-**g**, *Mtb*-induced oxidative burst in neutrophils exposed to the G6PD inhibitor G6PDi (**e**), the NOX2 inhibitor Diphenyleneiodonium chloride (DPI) (**f**), or PAD4 inhibitor Chlorine-amidine (Cl-amidine) (**g**) at 10, 50, and 100 ng/mL for 1 hour prior to infection. Column graphs (**e-g**) represent the magnitude of the oxidative burst and are shown as the mean ± SEM of the areas under the OCR curve for each group (n = 6-8 technical replicates per group). Statistical analysis: Data are presented as mean ± SEM; n = 5 biological replicates for metabolomics (a–c), and n = 6–8 technical replicates per group for oxidative burst assays (e–g). Statistical significance was determined using unpaired, non-parametric Mann–Whitney test for (a–c), and ordinary one-way ANOVA with Dunnett’s multiple comparisons test for (e–g) (\**P* < 0.03, \*\**P* < 0.002, \*\*\**P* < 0.0002, \*\*\*\**P* < 0.0001). All assays were performed independently three or more times.

To determine whether inhibiting metabolic flux through the PPP would abrogate the *Mtb*-induced neutrophil oxidative burst, we inhibited glucose-6-phosphate dehydrogenase (G6PD), the enzyme that catalyzes the first and rate-limiting step of the PPP by converting glucose-6-phosphate (G6P) to 6-phosphogluconolactone. G6PDi-1, a selective inhibitor of G6PD^78^, was added to neutrophil culture medium prior to *Mtb* infection. G6PDi-1 significantly reduced the *Mtb*-induced oxidative burst at all concentrations used (Fig. 7e). We next tested whether the *Mtb*-induced neutrophil oxidative burst could be suppressed by directly inhibiting NOX2. As shown in Fig. 7f, exposure of human neutrophils to diphenyleneiodonium chloride (DPI), a known inhibitor of NOX^79,80^, completely suppressed the *Mtb*-induced oxidative burst, consistent with findings from an unrelated study using PMA-stimulated neutrophils ^31^. Lastly, since NETosis can also occur through NOX2-independent pathways such as calcium ionophore or ATP mediated signaling^81^, we asked whether blocking NET formation via PAD4 inhibition would affect the *Mtb*-induced neutrophil oxidative burst. We exposed neutrophils to Cl-amidine, a PAD4 inhibitor^82^ shown to reduce bacterial burden, NETosis, and lung pathology in *Mtb*-infected mice^19^ and found that Cl-amidine significantly diminished the *Mtb*-induced oxidative burst (Fig. 7g).

Taken together, our findings demonstrate that *Mtb* infection triggers a profound metabolic reprogramming in human neutrophils, shifting glucose flux from glycolysis toward the PPP. This rerouting increases NADPH production, which fuels NOX2-dependent effector functions, including the oxidative burst and NETosis. Notably, these metabolic changes occur without major transcriptional alterations in canonical metabolic pathways, pointing to the integration of metabolic state with immune function in real time without requiring transcription. Pharmacological inhibition of G6PD (PPP flux), NOX2 (oxidative burst), or PAD4 (NETosis) markedly reduced *Mtb*-induced oxidative activity suggesting that *Mtb* drives neutrophils into a PPP-energized, NOX-2-mediated oxidative response that is tightly coupled to a PAD4-dependent NETosis machinery. Collectively, these findings define a PPP–NOX2–PAD4 immunometabolic axis that drives pathogenic neutrophil activity and highlight its components as potential targets for host directed therapies to limit neutrophil mediated tissue damage in TB.

## Discussion

In this study, we examined how *Mtb* infection reprograms the immunometabolism of human neutrophils in a fuel source-dependent manner, promoting enhanced oxidative burst and NETosis. Use of the Agilent XFe96 platform enabled high-resolution, real-time measurement of bioenergetic changes in *Mtb*-infected human neutrophils, allowing us to interrogate previously inaccessible aspects of their metabolic responses to infection. Notably, we found that *Mtb* induces a rapid and profound metabolic shift, redirecting glucose flux away from glycolysis and toward the PPP, thereby increasing NADPH production to fuel sustained NOX2-dependent ROS generation. These results, together with our findings that *Mtb* induces a neutrophil oxidative burst distinct from that triggered by NTMs, *Staphylococcus*, or *Listeria*, highlight a unique immunometabolic program in neutrophils that enables selective anti-*Mtb* effector functions during the early stages of infection. Importantly, our proteomic and mIF analyses provide clinically-relevant insights that cannot be obtained from animal models that do not fully replicate human TB pathology^10,35,36^. The significant enrichment of NETosis-associated proteins and GLUT3 in necrotic granulomas suggests that glucose utilization supports neutrophil activation within the human TB lesion microenvironment. Functional assays demonstrated that glucose availability governs neutrophil viability, degranulation, and oxidative responses, and that pharmacologic inhibition of G6PD, NOX2, or PAD4 suppresses the *Mtb*-induced oxidative burst. Unexpectedly, neutrophils exhibit remarkable metabolic plasticity during *Mtb* infection, maintaining cytokine production irrespective of fuel source even though glucose maximizes bioenergetic responses such as oxidative burst, OCR, and ECAR. Collectively, these results establish a mechanistic link between *Mtb*-driven metabolic reprogramming and neutrophil effector functions, identifying the PPP as a central metabolic driver of pathogenic neutrophil activity. Moreover, our data highlights GLUT3-dependent glucose transport and PPP flux as promising targets of HDT approaches to mitigate neutrophil-driven TB immunopathology.

NET components have been observed in the plasma of patients with TB^83^ and in TB lesions in mice^14^, but data linking nutrient transporters, metabolism, and neutrophil effector processes in human TB lesions has been lacking. Prior work showed that neutrophil activation increases reliance on extracellular glucose, largely through increased GLUT1 surface expression, and that glucose availability supports oxidative burst and NETosis^84^. However, these studies were performed in peripheral blood neutrophils and did not address spatial organization or tissue pathology. Similarly, studies in *Mtb*-infected nonhuman primates identified only GLUT1 as a hallmark of hypoxic macrophage zones surrounding necrotic cores^43^, leaving neutrophil-specific transporter usage unresolved. Our immunostaining revealed colocalization of GLUT3, but not GLUT1, with NET-associated proteins in human necrotic granulomas (Fig. 2). Since GLUT3 has a higher affinity for glucose than GLUT1^45,47^, this spatial association suggests a mechanistic link between high affinity glucose uptake, NETosis, and tissue necrosis in human TB granulomas. This represents a shift from prior models that emphasize GLUT1-dependent metabolism in NETosis during fibroblast inflammation^85^ and highlights GLUT3 as a potential regulator of neutrophil-mediated TB immunopathology. Consistent with the importance of glucose uptake for neutrophil effector function, genetic and pharmacologic inhibition of combined GLUT1 and GLUT3 dependent glucose transport has been shown to impair NADPH dependent oxidative burst in murine and human neutrophils during microbial challenge^86^. However, these studies did not resolve transporter specific functions within complex tissue environments. In contrast, our data show that in human necrotic granulomas, GLUT1 localizes predominantly outside neutrophil rich and NETosis active regions, while GLUT3 strongly colocalizes with neutrophil derived proteins including MPO, NE, and H3Cit (Fig. 2). Together, these results imply a division of labor in which GLUT1 supports macrophage metabolism⁴³, whereas GLUT3 fuels neutrophil driven NETosis, identifying GLUT3 as a dominant transporter associated with NET-mediated tissue destruction in human TB.

In parallel, we identified receptor-level mechanisms that drive the rapid oxidative burst in human neutrophils. *Mtb* induces a burst that is independent of TLR2/4 and Mincle, but is dependent, at least in part, on Dectin-1 and CD16, CD32, and CD64 (Fig. 5). This rapid, robust response distinguishes *Mtb* recognition from that elicited by NTMs, or other intracellular bacteria, and underscores pathogen-specific PRR signaling. These findings align with transcriptomic evidence of metabolically active CD64^high^ neutrophils as a permissive niche for *Mtb*^74^, and provide strong evidence that receptor-specific inputs drive the PPP-dependent oxidative program. Together, these findings identify Dectin-1 and FcγR signaling as important initiators of *Mtb*-induced oxidative responses with direct implications for HDT strategies. Consistent with this, recent work shows that mycobacteria exploit Dectin-1 signaling via a noncanonical α-glucan ligand to promote intracellular survival, underscoring the dual role of Dectin-1 in driving both host oxidative responses and pathogen persistence^63^.

Traditionally, HDTs against TB have targeted macrophage or CD8^+^ T cell metabolism^24,25^, but such approaches may not resolve neutrophil-driven pathology. Neutrophil depletion in mice reduces bacterial burden and lung pathology^16^, but carries the risk of severe bacterial and/or fungal infections given the central role of neutrophils in first line defense^18^. Importantly, neutrophils also engage in host-protective crosstalk with dendritic cells and T-cells during *Mtb* infection^87^, so indiscriminate depletion risks impairing adaptive responses and highlights the need for narrowly-focused targeting. Indeed, several studies have linked specific neutrophil activities to TB severity and immunopathology, including metabolically active pulmonary neutrophils that support *Mtb*^74^, type I IFN pathways that drive NET release and granuloma caseation^14,19^, and NOX2 pathway defects that expand immature, permissive neutrophils and lung inflammation^88^. Complementing these observations, inhibition of fatty acid oxidation in neutrophils has been shown to limit the accumulation of permissive immature subsets and reduce pathology and bacterial load *in vivo*, supporting metabolism-targeted intervention as a viable approach^89^. Our data refine this strategy by identifying neutrophil checkpoints tightly coupled to pathology, including restriction of PPP flux with G6PD inhibition, suppression of NOX2-mediated oxidative burst, and blockade of PAD4-dependent NET release, which has been shown to protect mice from severe TB immunopathology^19^. These interventions reduce the oxidative burst in our system while preserving core antimicrobial functions. These data, together with the localization of GLUT3, point to metabolic and receptor-specific pathways as practical therapeutic entry points for reducing neutrophil-driven tissue injury without neutrophil depletion. While this study identifies GLUT3, PPP activity, NOX2, PAD4, and receptor-mediated oxidative programs as potential HDT targets, the development of HDT strategies capable of reducing neutrophil-driven TB immunopathology will require systematic evaluation in experimental systems. Nonetheless, since animal models cannot reproduce the spatial and pathological heterogeneity of human TB, establishing the true clinical utility of neutrophil-targeted HDT regimens will require examination of human TB tissue.

In summary, our study uncovers *Mtb*-induced metabolic reprogramming as a central driver of neutrophil pathology, addressing a major gap in our understanding of human neutrophil metabolism in TB. Our data suggests that GLUT3 is a critical determinant of NETosis in necrotic human TB granulomas. Together with *Mtb*-induced PPP flux and receptor-driven rapid oxidative burst through Dectin-1 and FcγRs, these findings delineate key metabolic and signaling checkpoints that fuel damaging neutrophil effector responses. Importantly, we show that the oxidative burst induced by *Mtb* is rapid and pathogen-specific, underscoring unique host–pathogen interactions. These insights suggest promising HDT strategies that complement antibiotics by selective attenuation of destructive neutrophil functions while preserving antimicrobial defense. Collectively, our findings advance the understanding of neutrophil immunometabolism in TB and highlight actionable targets to limit immunopathology and improve clinical outcomes.

## Methods

### Human subjects

Approval for the ethical acquisition and examination of remnant human lung tissue specimens was obtained from the University of KwaZulu-Natal Biomedical Research Ethics Committee (BREC; approval numbers BCA 535/16 and BE019/13). Written informed consent was obtained from patients with TB undergoing lung resection at the King DinuZulu Hospital Complex, a tertiary center for TB treatment in Durban, South Africa. Collection of human blood was approved by the University of Alabama at Birmingham Institutional Review Board (IRB-300013604). Written informed consent was received from blood donors prior to their inclusion in this study. See Extended Data Table.1,3 for clinical details.

### Bacterial culture

*Mtb* H37Rv (BEI Resources, NR-123) or *Mtb* H37Rv expressing GFP (H37Rv-GFP) were used in all experiments. *Mtb* H37Rv-GFP was generated by transforming episomal plasmid pMV762^90^ encoding GFP into *Mtb* H37Rv followed by selection of positive colonies from Middlebrook 7H11 agar plates containing 50 µg/ml hygromycin B. Liquid cultures of *Mtb* H37Rv and H37Rv-GFP, *M. avium* 2285 Rough (BEI Resources, NR-44264) and *M. abscessus* (Moore and Frerichs) Kusunoki and Ezaki (ATCC, cat # 19977) were grown at 37 °C with shaking in Middlebrook 7H9 broth (Thermo Fisher cat # DF0713-17-9) supplemented with 0.2% glycerol, ADS (albumin [Millipore Sigma cat # 3116956001], dextrose [Thermo Fisher cat # DF0155-17-4], and sodium chloride), and 0.02 % tyloxapol (Sigma-Aldrich cat # T8761). H37Rv-GFP culture medium contained 50 µg/ml hygromycin B. Liquid cultures of *Staphylococcus aureus* SH1000 and *Listeria monocytogenes* 10403S (BEI cat # NR-13223) were grown at 37 °C with shaking in brain-heart infusion (BHI) broth.

### Neutrophil isolation

Blood was collected from healthy donors in vacutainers containing EDTA (BD, cat# 02-683-99C). Neutrophils were isolated by negative selection using an EasySep™ Direct Human Neutrophil Isolation Kit (Stemcell Technologies, cat # 19666) according to the manufacturer’s instructions. Isolated neutrophils were immediately used for experiments.

### Neutrophil viability assays

Freshly isolated neutrophils were infected with *Mtb* H37Rv at a multiplicity of infection (MOI) of 5 in RPMI 1640 medium containing 10 mM glucose, 1 mM HEPES, and 2 μg/mL Hoechst 33342 (Thermo Fisher cat # 62249). After 30 min, medium containing extracellular bacteria was removed and cells were washed three times with fresh RPMI 1640 medium containing 10 mM glucose and 1 mM HEPES. The wash medium was then replaced with RPMI 1640 containing various nutrients at 10 mM final concentration, 1 mM HEPES, and 0.5 μM propidium iodide (Thermo Fisher cat # P1304MP). Neutrophil viability was monitored over 24 h by quantifying PI-positive cells at 1 h intervals using an Agilent BioTek Cytation 5 Cell Imaging Multimode Reader.

### Histology and multiplex immunofluorescence microscopy

5 µm-thick sections were cut from formalin-fixed, paraffin-embedded (FFPE) human TB lung tissues, and a single tissue section was used for H&E staining, DAPI staining, and mIF microscopy essentially as described^4^. Automated sequential immunofluorescence staining of human lung tissue sections was performed using the Lunaphore COMET Platform^91^. Tissue sections were analyzed using the following primary antibodies: MPO (Invitrogen cat # MA1-80878; 1:200), Cit-Histone H3 (Invitrogen cat # 630-180ABBOMAX; 1:200), NE (Invitrogen cat # MA5-42901; 1:1000), GLUT-1 (Invitrogen cat # MA5-31960; 1:10,000), and GLUT-3 (Invitrogen cat # MA5-32697; 1:500). For each iterative round of staining, corresponding secondary antibodies conjugated to Cy5 (Proteintech CoraLite Plus 647 cat # RGAR005; 1:200) or TRITC (Proteintech CoraLite Plus 555 cat # RGAR003; 1:200) were applied. DAPI (Thermo Fisher, cat # 62248, 1:1000) was used as a nuclear counterstain. Post-processing analysis included subtraction of endogenous tissue autofluorescence to improve visualization. Image analysis and figure generation were performed using HORIZON software (v.2.1.0.0).

### Laser-capture microdissection and protein extraction

FFPE human TB lung tissues were cut into 30 µm-thick sections and mounted on membrane-coated slides. Regions of interest (ROI) were excised and collected by laser capture microdissection (LCM) using a Nikon Ti Eclipse microscope equipped with the MMI CellCut system, controlled through MMI software, and fitted with a Nikon Digital Sight DS-U2 camera for imaging. Each ROI was transferred to a well of a protein LoBind 96-well plate (Eppendorf, cat# 0030129504) containing 50 µL of lysis buffer consisting of 50 mM triethylammonium bicarbonate (TEAB; Thermo Fisher, cat# 90114) and 5% SDS (Thermo Scientific, cat# 15553027). Samples were sonicated using a BeatBox™ (PreOmics GmbH) at high power for 10 minutes, briefly centrifuged, and heated at 90 °C for 1 hour for protein denaturation. The plate was sonicated again for an additional 10-minute high-power cycle, followed by centrifugation at 3,000 rpm. Protein concentration was determined using a Pierce™ BCA Protein Assay kit (Thermo Scientific, 23225).

### Sample preparation for LC–MS/MS

Proteins were reduced by adding 0.5 µL of 500 mM Tris (2-carboxyethyl) phosphine hydrochloride (TCEP; Sigma-Aldrich, C4706) to a final concentration of 5 mM and incubating samples at 56 °C for 15 minutes. Alkylation was performed by adding 2 µL of 500 mM chloroacetamide (CAA; Millipore) to a final concentration of 20 mM, followed by incubation for 30 minutes in the dark. Samples were acidified by adding 5 µL of 27.5% aqueous phosphoric acid to a final concentration of 1.6%. Protein digestion and cleanup were performed using the S-Trap™ 96-well MS Sample Prep Kit (Protifi, NC1508276) according to the manufacturer’s protocol. Briefly, 350 µL of S-Trap binding buffer (90% methanol, 100 mM TEAB, pH 7.1) was added to the acidified lysate, mixed thoroughly, and transferred to the S-Trap plate positioned atop a 96-well receiver plate, followed by centrifugation at 1,500 g for 2 minutes. As recommended for FFPE tissues, a chloroform wash was performed using 150 µL of 50% methanol/50% chloroform, followed by centrifugation at 1,500 g for 2 minutes. Captured proteins were washed three times with 200 µL of S-Trap binding buffer. For on-column digestion, the S-Trap plate was transferred to a clean receiver plate, and 125 µL of digestion buffer containing TPCK-treated trypsin (Thermo Scientific, 20233) at a 1:10 enzyme-to-protein ratio was added. Samples were incubated overnight at 37 °C. Peptides were sequentially eluted by centrifugation following the addition of 80 µL of digestion buffer, 80 µL of 0.2% aqueous formic acid, and 80 µL of 50% aqueous acetonitrile containing 0.2% formic acid, resulting in a final acetonitrile concentration of approximately 10% (v/v). Eluted peptides were dried using a vacuum centrifuge. Dried peptides were resuspended in 0.1% aqueous formic acid, and peptide concentration was determined using a Pierce™ Quantitative Fluorometric Peptide Assay kit (Thermo Scientific, 23290). Peptides were further desalted using Pierce™ C18 Spin Columns (Thermo Scientific, 89870) according to the manufacturer’s instructions. Columns were activated twice with 200 µL of 50% methanol and equilibrated twice with 200 µL of 0.5% trifluoroacetic acid (TFA) in 5% acetonitrile (ACN). Samples were loaded onto the columns, with the flow-through reapplied once to maximize binding, washed twice with 0.5% TFA in 5% ACN, and eluted twice with 20 µL of 70% ACN. Eluted peptides were dried using a vacuum evaporator, reconstituted in 30 µL of 2% ACN containing 0.2% formic acid, and diluted to a final concentration of 100 ng/µL for LC–MS/MS analysis.

### Liquid chromatography-tandem mass spectrometry

For proteomic analysis, 200 ng of peptides per sample were separated on an Aurora Ultimate UHPLC column (25 cm × 75 µm, 1.7 µm C18; IonOpticks) maintained at 50 °C. Samples were loaded directly onto the analytical column without a trapping column to maximize sensitivity. Peptide separation was performed on a Vanquish Neo UHPLC system (Thermo Fisher) coupled to an Orbitrap Exploris 480 mass spectrometer (Thermo Fisher) equipped with a Nanospray Flex ion source, using a 123-minute linear gradient at a flow rate of 0.22 µL/min.

Solvent A consisted of 0.2% formic acid in 2% acetonitrile, and solvent B consisted of 0.2% formic acid in 80% acetonitrile. The gradient was initiated at 3% solvent B and held for 7 minutes, followed by a linear increase to 25% solvent B over 83 minutes. Solvent B was then increased to 41% over the next 32 minutes, followed by a rapid ramp to 95% solvent B over 1 minute for column washing. The column was subsequently re-equilibrated to initial conditions prior to the next injection. Data-dependent acquisition was performed in positive ion mode with a spray voltage of 1,600 V and an ion transfer tube temperature of 300 °C. Full MS scans were acquired at a resolution of 60,000 over an m/z range of 375–1,200, with a 3-second cycle time, an AGC target of 300%, and automatic injection time. Precursor ions with charge states of +2 to +6 and intensities above 5 × 10³ were selected for fragmentation, with dynamic exclusion applied after a single acquisition (45 seconds, 10 ppm). MS/MS spectra were acquired in the Orbitrap at 30,000 resolution using a 1.6 m/z isolation window, higher-energy collisional dissociation (HCD) with a normalized collision energy of 28%, an AGC target of 200%, and automatic injection time. Data were acquired using Xcalibur software (Thermo Scientific).

### LC-MS/MS data analysis and statistics

LC–MS/MS data were acquired in data-dependent acquisition (DDA) mode and processed using the FragPipe computational pipeline. Raw mass spectrometry files were analyzed with FragPipe v21.1^92,93^, incorporating MSFragger v3.8^94^ for database searching and Philosopher v5.1.1^95^ for downstream processing. Spectra were searched against the UniProt human reference proteome (20,631 entries) along with *Mycobacterium tuberculosis* proteomes (strains ATCC 25177, ATCC 25618, ATCC 35801, ATCC 35801, CDC 1551, and K85 totaling 25407 protein sequence entries) with trypsin as the proteolytic enzyme, allowing up to two missed cleavages. Carbamidomethylation of cysteine residues was specified as a fixed modification, while oxidation of methionine and acetylation of protein N-termini were considered variable modifications. The precursor mass tolerance was set to ±20 ppm for both initial and main searches, and the fragment mass tolerance was set to 20 ppm, with peptide lengths restricted to 7–50 amino acids. False discovery rates for peptide-spectrum matches, peptides, and proteins were controlled at 5% using the target–decoy strategy implemented in Philosopher.

Label-free protein quantification was performed using IonQuant v1.10.11^96^ within FragPipe. Quantification was based on precursor-level extracted ion chromatograms with a mass tolerance of 10 ppm, retention time tolerance of 0.4 min, and isotope mass tolerance of 20 ppm. Protein abundances were estimated using the top three peptide intensities per protein, and global intensity-based normalization was applied to reduce systematic variation across samples. Processed quantitative protein tables were subsequently imported into the tidyproteomics R package (v1.8.8)^97^ for statistical and exploratory analysis. Data were filtered to retain proteins identified by at least two unique peptides and quantified in the majority of samples per condition. Intensities were log₂-transformed and normalized using an SVM regression to correct for remaining technical variation. Principal component analysis and hierarchical clustering were used to assess sample-level variation and reproducibility. Differential protein abundance was evaluated Student’s t-test^98^, and p-values were adjusted for multiple testing using the Benjamini–Hochberg^99^ procedure to control the false discovery rate.

### NE, MPO, and NETosis assays

Freshly isolated human neutrophils were infected with *Mtb* H37Rv at a multiplicity of infection (MOI) of 5 in RPMI 1640 containing 10 mM glucose and 1 mM HEPES. After 30 min, medium containing extracellular bacteria was removed and cells were washed three times with glucose-free RPMI 1640 containing 1 mM HEPES. After removing the wash buffer, infected neutrophils were cultured for 2 hours in RPMI 1640 containing various nutrients at a final concentration of 10 mM. Quantitation of MPO and NE in the culture medium was performed by measuring MPO activity (Cayman Chemicals, cat # 600620) or NE activity (Cayman Chemicals, cat # 600610 according to the manufacturer’s protocols. NETosis was visualized by staining nuclear and extruded genomic DNA (Cayman Chemicals, cat # 601750) according to the manufacturer’s instructions. Absorbance measurements for MPO and NE activity and NETosis visualization were performed using an Agilent BioTek Cytation 5 Cell Imaging Multimode Reader. Image analysis was performed with Agilent BioTek Gen5 software.

### Neutrophil extracellular flux analysis

The extracellular acidification rate (ECAR), neutrophil oxidative burst and mitochondrial respiration [oxygen consumption rate (OCR)] were measured using the Agilent Seahorse XFe96 Extracellular Flux Analyzer with XFe96 FluxPaks (Agilent cat # 102416-100). All other reagents used were purchased from Millipore Sigma. Human neutrophils were isolated as described above and used immediately for assays. For each experiment, 1 × 10^5^ cells per well were seeded into poly-D-lysine-coated (Thermo Fisher cat # A3890401) XFe96 culture plates.

The oxidative burst following PMA stimulation was measured by injecting PMA (100 ng/mL final concentration) through port A. For infection assays, neutrophils were first exposed to *Mtb* H37Rv at an MOI of 5 for 30 min, washed three times with glucose-free RPMI 1640, and then cultured in RPMI 1640 containing various nutrients (10mM final concentration). In a modified assay, rotenone (2.5 µM final concentration) and antimycin A (1.25 µM final concentration) were injected first (port A) to inhibit mitochondrial oxygen consumption, followed by injection of either PMA or bacteria (MOI 5) through port B, and in some experiments, glucose (10 mM final concentration) was injected third through port C. For oxidative burst experiments involving pretreatments, neutrophils were incubated for 30 min with *Mtb* fractions, recombinant cytokines, inhibitors, or blocking antibodies prior to measuring the oxidative burst. Final working concentrations of all blocking reagents are provided in the corresponding figure legends. All *Mtb* cell fractions were obtained from the BEI Resources Repository (Cell Wall, cat # NR-14828; Whole Cell Lysate, cat # NR-14822; Total Lipids, cat # NR-14837; Cell Membrane, cat # NR-14831) and were used at a final concentration of 25 µg/ml. Irradiated H37Rv (BEI cat # NR-49098) was used at an MOI of 5. Blocking antibodies were sourced as follows: Mincle (Invivogen cat # mabg-hmcl-2), Dectin-1 (Invivogen, cat # mab9-hdect1), CD16 (BioLegend cat # 302001), CD32 (StemCell Technologies, cat # 60012), and CD64 (BioLegend cat # 305002). TLR inhibitors TLR2-IN-C29 (cat # SML3817) and TAK-242 (cat # 614316) were purchased from Millipore Sigma. Recombinant cytokines were obtained from BioLegend: human IFN-γ (cat # 570204), mouse GM-CSF (cat # 576304), and human TNF-α (cat # 570102). Additional inhibitors were purchased from Cayman Chemical: g6PDi-1 (cat # 31484), diphenyleneiodonium (cat # 81050), and Cl-amidine (cat # 10599). Oxidative burst was quantified as the area under the curve (AUC).

Neutrophil ECAR and OCR assays were performed as described by us and others^26,53^, with modifications. Unlike macrophages, which were infected with *Mtb* prior to assay^26^, neutrophils were exposed to bacteria via injection through port A together with 10 mM glucose, followed by second injection of oligomycin (2.5 µM final concentration), third injection of FCCP (1.5 µM final concentration) and pyruvate (1 mM final concentration) and final injection of rotenone (2.5 µM final concentration) and antimycin A (1.25 µM final concentration). All data were normalized to 1x10^4^ cells by staining nuclei with Hoechst 33342 (2 µg/mL; Thermo Fisher cat # 62249) and counting cells using an Agilent BioTek Cytation 5 Cell Imaging Multimode Reader with Gen5 software.

### Metabolomics

Neutrophils (3 × 10^6^ cells/well) were seeded into poly-D-lysine–coated six-well plates and infected with *Mtb* at an MOI of 5 for 30 min in RPMI 1640 containing 10 mM unlabeled glucose. Following infection, cells were washed three times with glucose-free RPMI 1640 and incubated at 37 °C for 30 min in glucose-free RPMI 1640. Cells were then washed again glucose-free RPMI and cultured for 30 min in RPMI 1640 containing 10 mM ^13^C_1,2_-glucose for metabolic incorporation. After incorporation, the medium was removed, and cells were washed with warm PBS. Metabolites were extracted in an ice-cold 1:1 mixture of methanol and water. Deuterated succinic acid-2,2,3,3-d_4_ (Millipore Sigma cat# 293075) was included at the time of quenching as an internal standard (10 ng/mL). Extracts were subjected to three freeze-thaw cycles, centrifuged through 0.22 μm spin filters, and protein concentration was determined using a BCA assay. Samples were dried under vacuum and stored at -80 °C until analysis. For LC–MS/MS, dried metabolites were reconstituted in 150 μL water, passed through 0.22 μm filters, and analyzed. Commercial standards (Millipore Sigma) for all metabolites were used to validate peak identification. Relative metabolite abundances were normalized to internal standards (deuterated succinate) and to protein content. Metabolite quantitation, data processing, and visualization were performed using Skyline (v4.1.0; MacCoss Lab Software) and Metaboanalyst 6.0.

### Cytokine quantitation

Freshly isolated neutrophils from healthy donors were cultured in serum-free RPMI 1640 containing various carbon sources at 10 mM. Neutrophils were infected with *Mtb* H37Rv at an MOI of 5 for 1 hr at 37 °C. After 1 hr, medium from infected and uninfected controls was collected, cleared by centrifugation, and passed through a 0.22 μM spin filter to ensure sterility. Cytokine quantitation was performed using a Bio-Plex Pro Human Cytokine 17-plex Assay kit (Bio-Rad cat # M5000031YV) following the manufacturer’s protocol. Briefly, medium samples were incubated with magnetic beads coated with capture antibodies on a shaker at 850 rpm for 30 min at RT, washed twice in Bio-Plex wash buffer, and incubated with biotinylated detection antibodies. Streptavidin conjugated to phycoerythrin (PE) was then added, followed by three final washes. Beads were resuspended in assay buffer, and fluorescence was measured on a MAGPIX® instrument (Luminex Corp). The readouts were analyzed with the standard version of EMD Millipore’s Milliplex® Analyst software (Millipore-Sigma and VigeneTech, Inc.) and the cytokine concentrations were determined using a fluorescence standard curve generated from assay standards.

### RNA-Seq gene expression analysis

RNA was isolated from freshly isolated neutrophils infected with *Mtb* H37Rv at an MOI of 5 for 1 hr and uninfected controls. Cells were lysed and RNA extracted using a NucleoSpin RNA kit (Takara Bio USA Inc., cat # 740955) following the manufacturer’s protocol. RNA quality and integrity were assessed using the Agilent 2100 Bioanalyzer, and only samples with RIN ≥ 7.0 were used for library preparation. Sequencing libraries were prepared using a NEBNext Ultra II Directional RNA Library Prep Kit with poly(A) selection (New England Biolabs cat # E7760L) according to the manufacturer’s recommendations. Libraries were quantified by qPCR using the Kapa Biosystems Library Quantification Kit (Roche LightCycler 480) before cluster generation. High-throughput sequencing was carried out on the Illumina NextSeq 500 platform using NextSeq software v4.2.0, generating paired-end reads. Raw FASTQ files were aligned to the GRCh38 reference genome (GENCODE Release M24) with STAR (v2.7.3a). Read counts were assigned to genes using HTSeq-count (v0.13.5), and differential gene expression analysis was performed using DESeq2, applying the Benjamini–Hochberg procedure to control the false discovery rate.

### Glucose quantitation

Glucose content in lung tissue was measured using an Abcam Glucose Assay Kit, fluorometric format (cat # ab65333). Approximately 10 mg of tissue was weighed and homogenized in assay buffer using a handheld Dounce homogenizer, and insoluble debris was removed by centrifugation. To eliminate protein interference, tissue supernatants were deproteinized by treatment with 1M perchloric acid, followed by neutralization with 2M potassium hydroxide, ensuring a final pH of 6.5–8. The clarified extracts were diluted four-fold in assay buffer to fall within the linear range of the glucose standard curve. Samples were incubated with the assay reaction mix for 30 min at 37 °C in the dark and fluorescence (ex/em = 535/587 nm) was measured using a Hidex plate reader within BSL3 containment. Reader settings were optimized to minimize background, and each plate was read 5–6 times, with glucose concentrations calculated independently from each replicate readout. Concentrations were normalized to the tissue weight. For comparison between granulomatous and healthy control tissues, replicate values per sample were averaged, and mean concentrations were reported for each group.

### Statistical Analysis

All experiments were performed at least three times independently with 5-8 technical replicates. For metabolomics and transcriptomics, five and three independent biological replicates were used, respectively. Statistical analyses were performed using GraphPad Prism 10 (GraphPad Software, Inc., La Jolla, USA). Statistical significance was defined as: \**p*<0.03, \*\**p*<0.002, \*\*\**p*<0.0002, \*\*\*\**p*<0.0001. Statistical tests used in each figure are described in the figure legend.

## Data availability

All source data will be made available with the publication of the manuscript.

## Author contributions

Conceptualization: KCC, HTP, AJCS, AA. Methodology: KCC, AJCS. Formal analysis: KCC. Investigation: KCC, HTP, DN, SN, RS, VP, TN, GW, KL, KN, AG, JJJ, TFC, MP, MLR, JNG. Proteomics: MP, TY, BQ, JJJ. Proteomics

Analysis: JJJ. Visualization: KCC, AJCS. Writing – Original Draft: KCC. Writing – Review and Editing: KCC, JNG, AJCS. Supervision: JNG and AJCS. Funding Acquisition: AJCS.

## Acknowledgements

This work was supported by NIH grants R01Al111940, R01AI134810, R01AI137043, R33AI138280, R21A127182, and pilot funds from the UAB CFAR and UAB Heersink School of Medicine to AJCS. Additional support was received from the Global Research Resource for Human Tuberculosis (NIH R24AI186591 to AJCS) and the UAB-UCSD O’Brien Core Center for Acute Kidney Injury Research (NIH U54 DK137307 to AA). The authors thank Dr. Sixto Leal and Dr. Sawanan Saitornuang (UAB) and Dr. Lorenzo Thompson (UAB Immunology Institute) for assistance with human blood collection, and Dr. Megan Kiedrowski (UAB) for generously providing *Staphylococcus aureus* SH1000. The authors would also like to thank Dr. Davide Botta and the Antibody Characterization and Serology facility of the UAB Immunology Institute for performing and analyzing multiplex immunoassays. The authors thank Dr. Tsui-Fen Chen (Caltech) for her assistance with proteomics. This work was supported by the staff and resources of the UAB Southeastern Biosafety Laboratory Alabama Birmingham (SEBLAB), a NIAID-supported (UC6 AI058599, G20 AI058599, and UC7 AI180255) Regional Biocontainment Laboratory.

## Notes

### Competing Interest Statement

The authors have declared no competing interest.

