## Extended Data and Tables for "Glucose selectively drives a rapid oxidative burst and immunometabolic reprogramming in human neutrophils during *Mycobacterium tuberculosis* infection"

Extended Data Figures and Tables

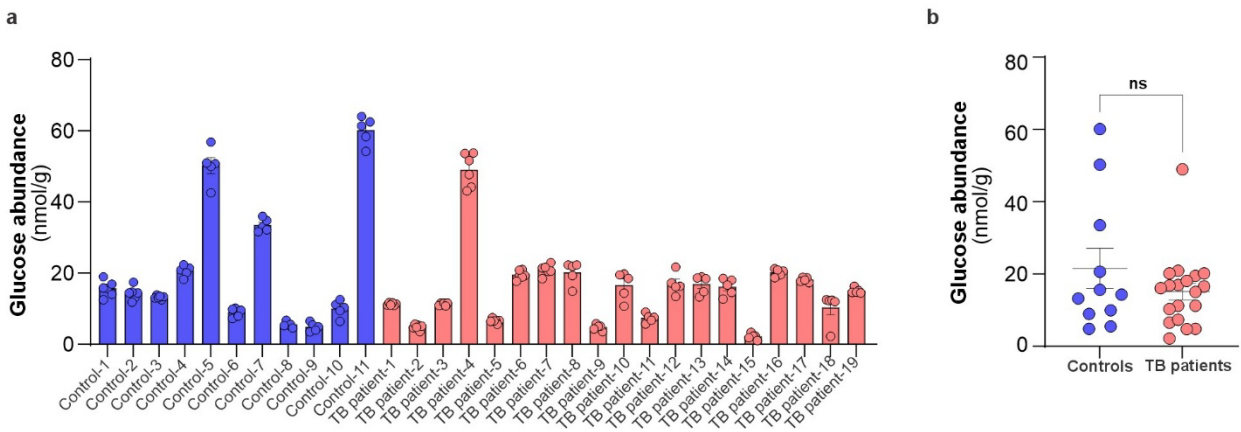

**Extended Data Fig. 1 | Glucose abundance in human necrotic granulomas.** **a**, Glucose abundance was measured in human necrotic granulomas from 19 TB patients and in 11 healthy control lung tissue specimens from non-TB patients. **b**, Statistical analysis shows no significant difference in glucose abundance between TB granulomas and control specimens. Glucose concentration is normalized to tissue weight, and the statistical significance was calculated using the nonparametric Mann–Whitney t-test, $P = 0.703$.

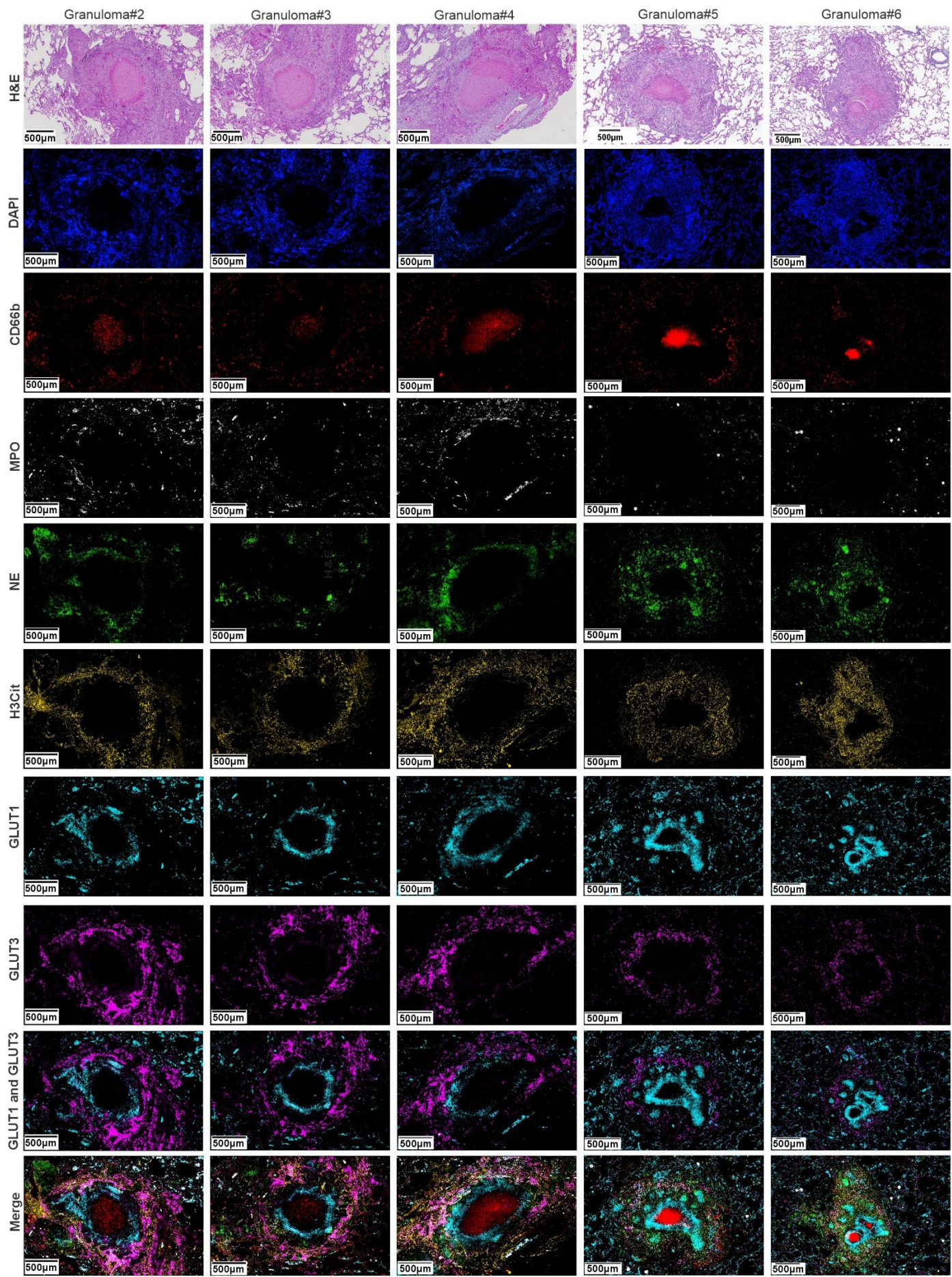

**Extended Data Fig. 2 | Colocalization of GLUT3 with NETosis markers in human TB granulomas.** H&E and DAPI staining and multiplex immunofluorescence (mIF) microscopy was performed on five representative necrotic TB granulomas from two patients with TB. The distribution of neutrophil/NETosis markers CD66b, MPO, NE, and H3Cit, and GLUT1 and GLUT3 is shown. Merged images show localization of GLUT3 with NETosis markers. Autofluorescence subtraction and image processing were performed using HORIZON software.

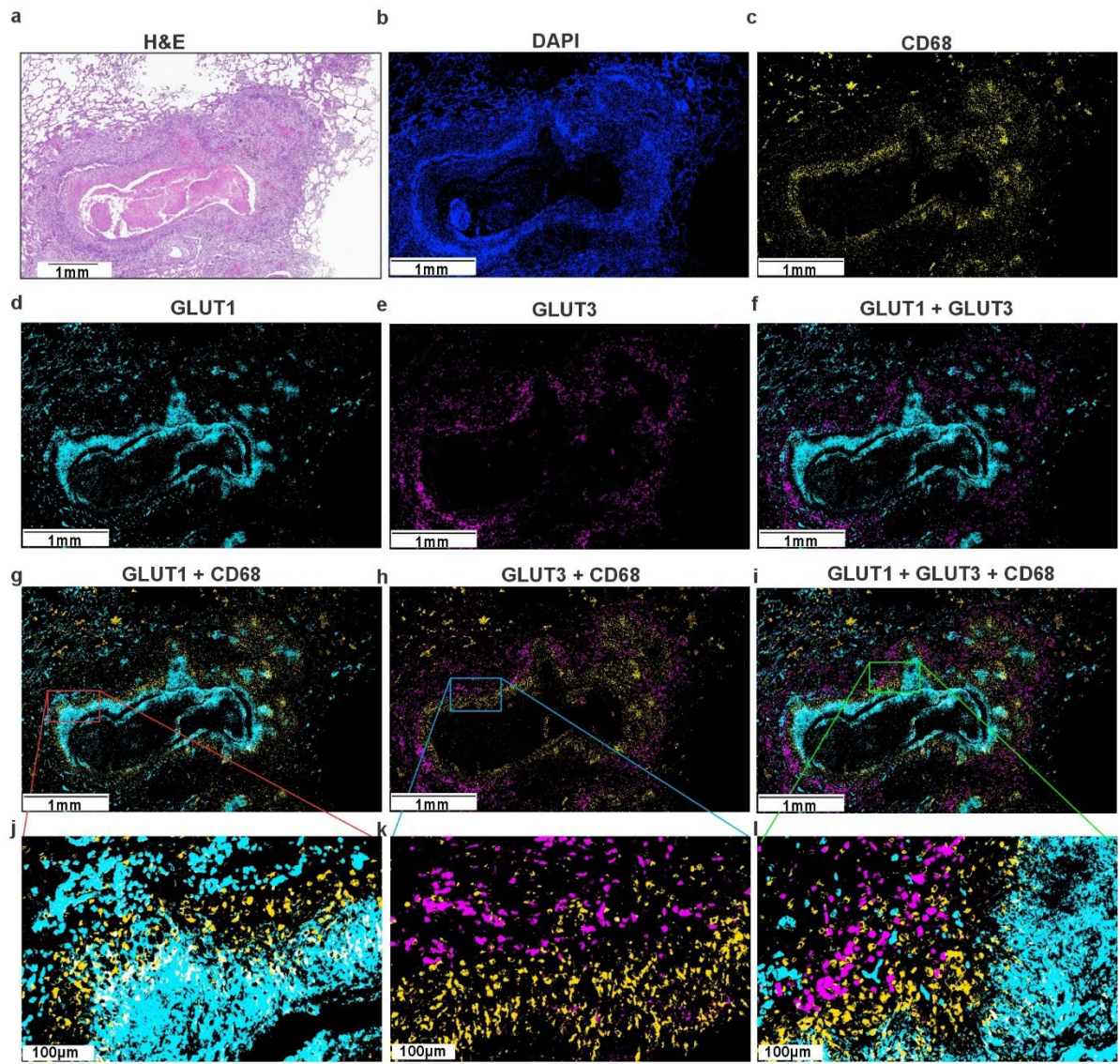

**Extended Data Fig. 3 | Macrophage-defined separation of GLUT1- and GLUT3-positive metabolic zones in human tuberculosis granulomas.** **a**, H&E and **b**, DAPI staining of a representative necrotic TB granuloma from a single TB patient section. **c–i**, Multiplex immunofluorescence (mIF) microscopy identifies cell type and metabolic marker distribution. Macrophages were identified by CD68 expression (**c**). GLUT1 (**d**) and GLUT3 (**e**) expression was detected in distinct pathological regions of the granuloma. **g,j**, Merged images of GLUT1 and CD68 show partial colocalization of CD68<sup>+</sup> macrophages with GLUT1 in the peri-necrotic region. **h,k**, Minimal colocalization was observed between CD68<sup>+</sup> macrophages and GLUT3. **i,l**, CD68<sup>+</sup>, GLUT-negative macrophages form an intermediate ring between the GLUT1<sup>+</sup> inner region and the peripheral GLUT3<sup>+</sup> zone. Autofluorescence subtraction and image processing were performed using HORIZON software.

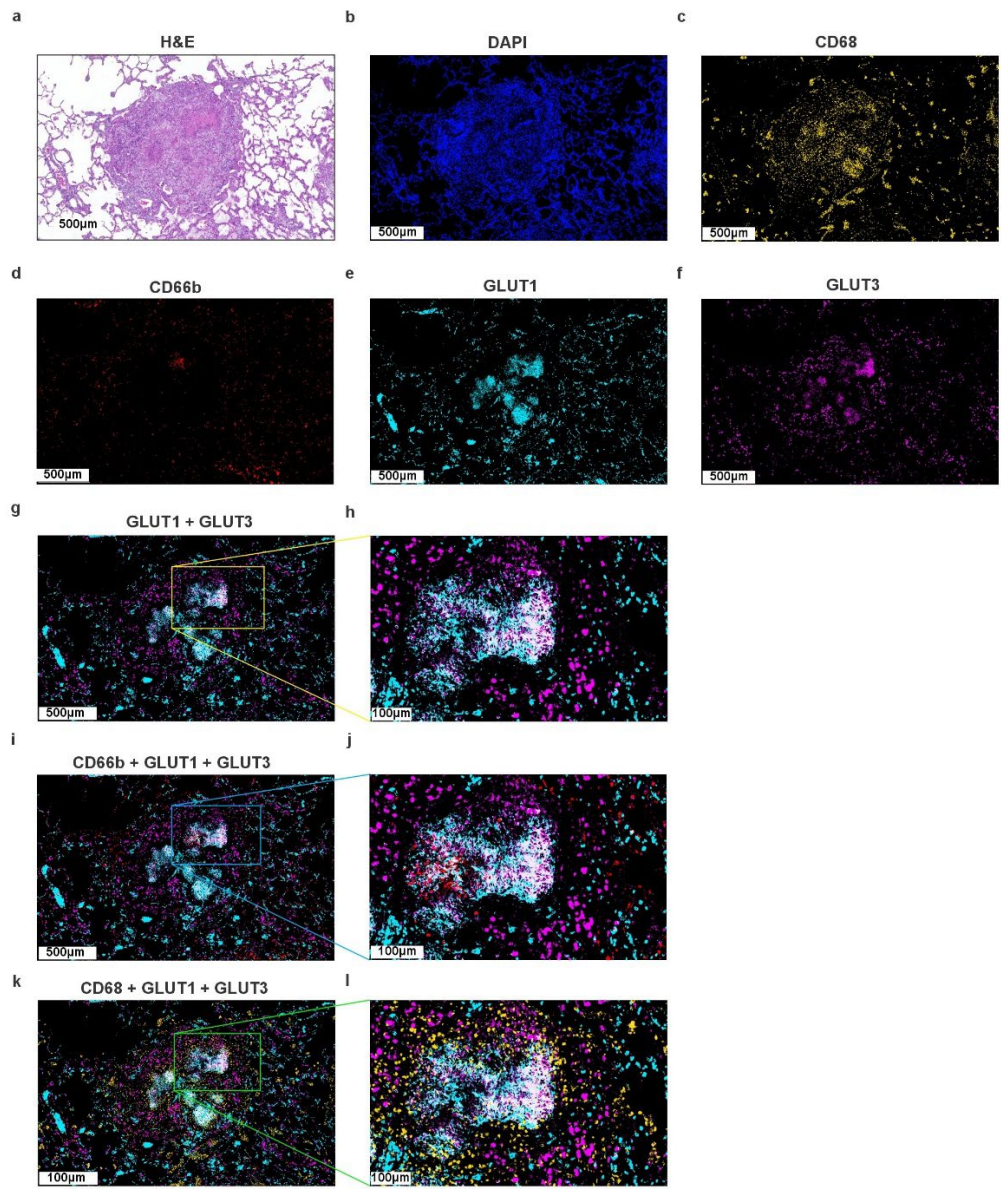

**Extended Data Fig. 4 | GLUT1 and GLUT3 exhibit overlapping spatial distribution in non-necrotic human TB granulomas.** (a) H&E and (b) DAPI staining of a representative non-necrotic TB granuloma from a single patient section, showing little, if any, necrotic core. (c–l) Multiplex immunofluorescence (mIF) microscopy was performed to determine the distribution of cell types and metabolic markers. (c) Macrophages were identified by CD68 expression. (d) CD66b<sup>+</sup> neutrophils were dispersed throughout the granuloma with positive DAPI staining, and no central CD66b<sup>+</sup> necrotic core was observed. (e) GLUT1 and (f) GLUT3 expression was detected across the non-necrotic granuloma. (g,h) Merged images of GLUT1 and GLUT3 show significant colocalization (white region). (i,j) CD66b<sup>+</sup> neutrophils were present within regions co-expressing GLUT1 and GLUT3. (k,l) CD68<sup>+</sup> macrophages were also present within GLUT1<sup>+</sup>/GLUT3<sup>+</sup> regions. Autofluorescence subtraction and image processing were performed using HORIZON software.

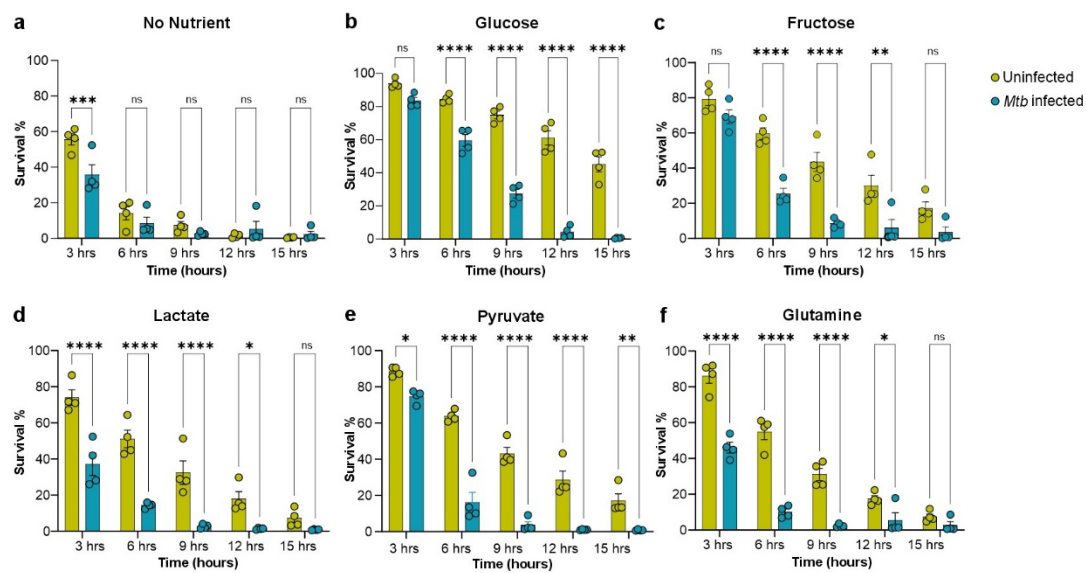

**Extended Data Fig.5 | *Mtb* infection decreases the viability of human neutrophils.** Survival of uninfected and *Mtb*-infected (MOI of 5) neutrophils cultured in with (a) no nutrient or (b) glucose (c) fructose (d) lactate (e) pyruvate (f) glutamine at 10 mM. Column graphs (from viability curves in Fig. 3) show neutrophil viability at 3, 6, 9, 12, and 15 hours with and without infection. Data are shown as mean ± SEM; $n = 4$ technical replicates per group. Statistical significance was determined using two-way ANOVA with Tukey's multiple comparisons test (\*$P < 0.03$, \*\*$P < 0.002$, \*\*\*$P < 0.0002$, \*\*\*\*$P < 0.0001$).

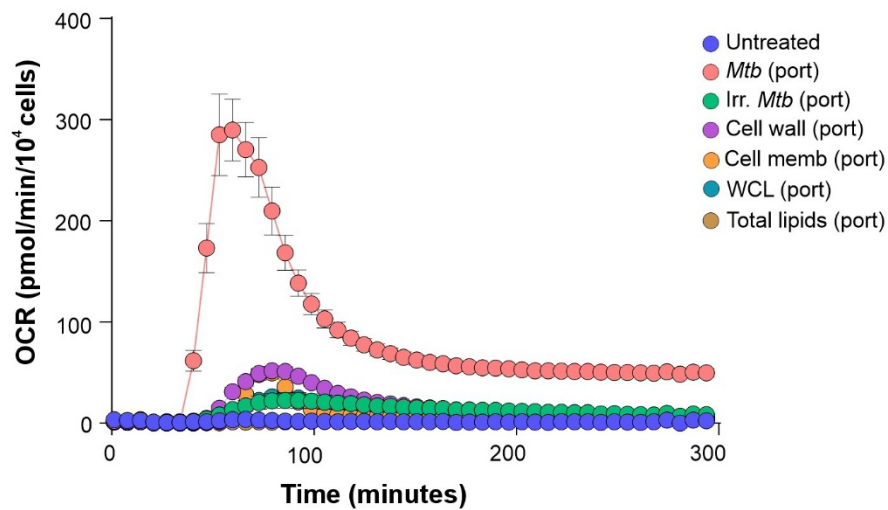

**Extended Data Fig. 6 | Live *Mtb* induces maximal neutrophil oxidative burst compared to irradiated bacteria or isolated cellular components.** (related to Fig. 4g). Oxidative burst of neutrophils cultured on glucose (10 mM) in response to live *Mtb* (MOI of 5), irradiated (Irr.) *Mtb*, *Mtb* whole-cell lysate (WCL), *Mtb* cell wall, *Mtb* cell membrane, or total *Mtb* lipid fractions.

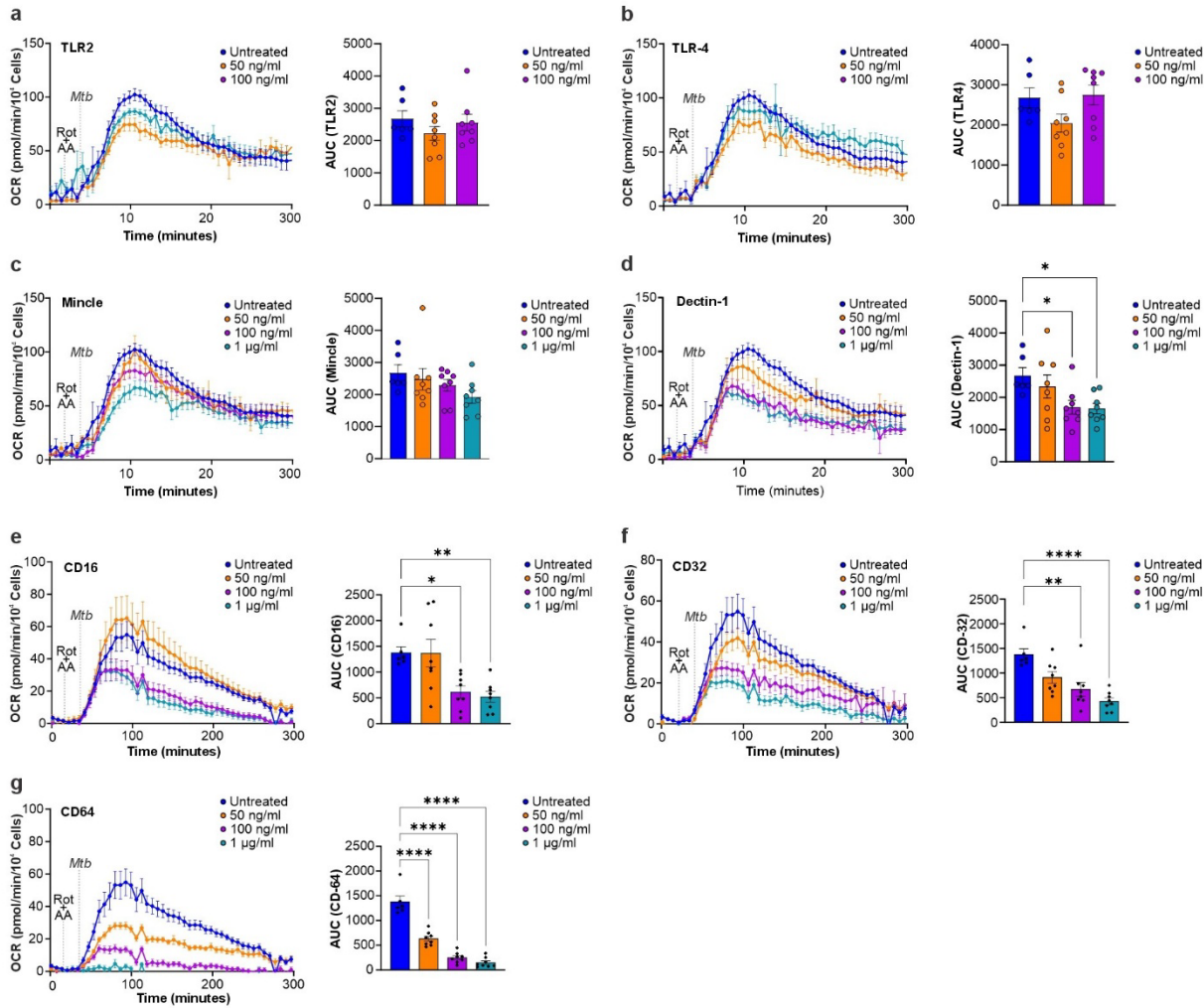

**Extended Data Fig. 7 | *Mtb*-induced oxidative burst is inhibited by receptor-specific blockade in human neutrophils.** **a-g**, The oxidative burst in neutrophils from healthy donors was measured by extracellular flux analysis and quantified as area under the OCR curve (AUC). All XF data were normalized to 10,000 cells using Gen5 software on an Agilent BioTek Cytation 5 Multimode Reader. Serum-free RPMI 1640 medium was used in all assays. One hour prior the start of the assay, neutrophils were exposed to the TLR2 inhibitor TLR2-IN-C29 at 50 and 100ng/mL (**a**) or TLR4 inhibitor TAK-242 at 50 and 100ng/mL (**b**) or blocking antibodies directed against Mincle (**c**), Dectin-1 (**d**), CD16 (**e**), CD32 (**f**), or CD64 (**g**) at 50 ng/mL, 100ng/mL, and 1 µg/mL. Rotenone (1.25 µM) and antimycin A (2.5 µM) were then injected via port A to inhibit mitochondrial respiration, followed by *Mtb* injection via port B (MOI of 5). Column graphs represent the magnitude of the oxidative burst are shown as the mean ± SEM; n = 6–8 per group; ordinary one-way ANOVA with Dunnett's test and statistical significance shown as \*$P < 0.03$, \*\*$P < 0.002$, \*\*\*$P < 0.0002$, \*\*\*\*$P < 0.0001$.

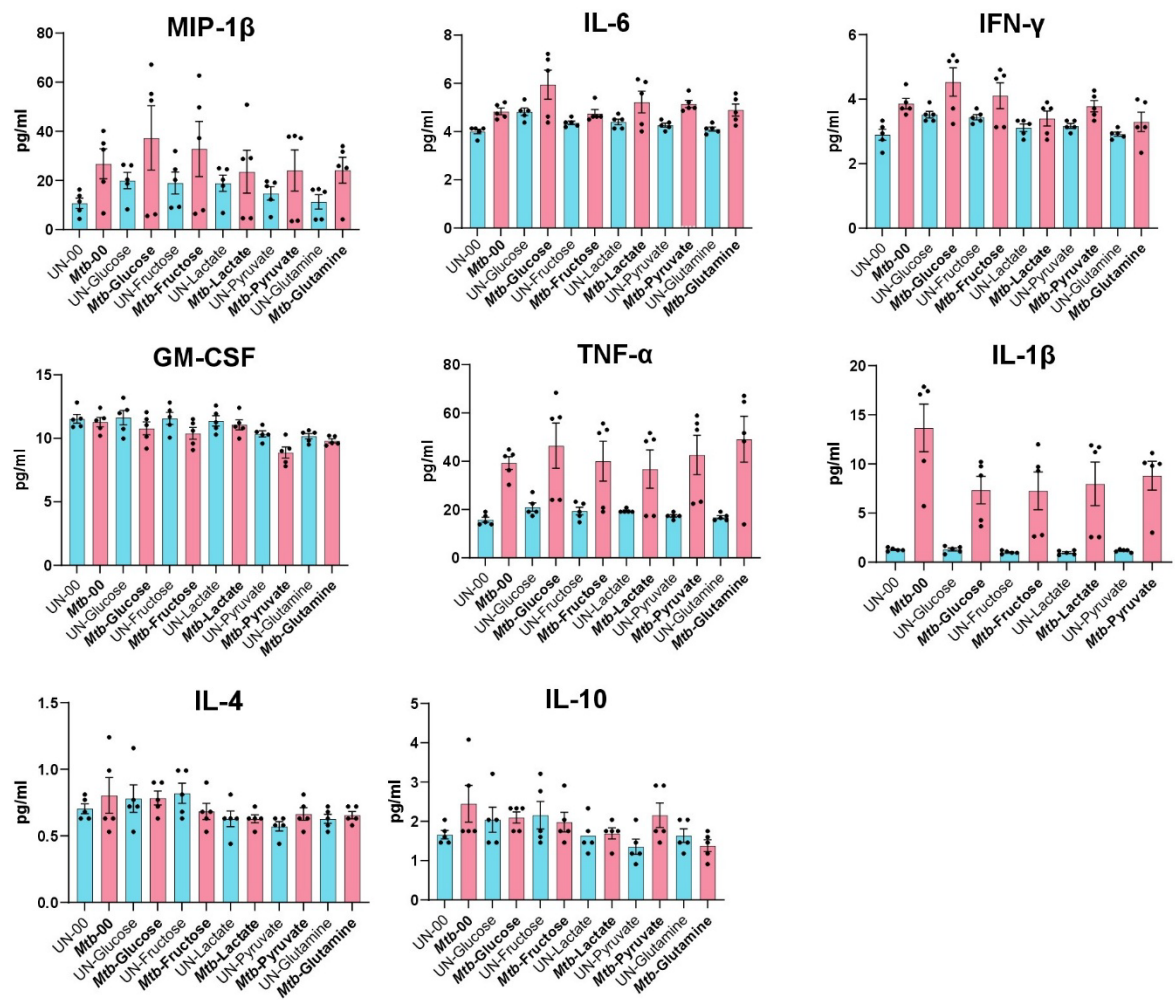

**Extended Data Fig. 8 | Cytokine secretion by human neutrophils upon *Mtb* infection.** Freshly isolated neutrophils from healthy donors were cultured in serum-free RPMI 1640 medium supplemented with individual carbon sources (10 mM each). Cells were infected with *Mycobacterium tuberculosis* (*Mtb*) H37Rv at a multiplicity of infection (MOI) of 5 for 1 h. Culture supernatants were collected, clarified by centrifugation, sterile-filtered, and subjected to multiplex cytokine profiling. Data represent mean ± SEM from 5 independent biological replicates. Cytokine differences across nutrient conditions were analyzed using ordinary one-way ANOVA with Dunnett's post hoc multiple-comparisons test against the control condition.

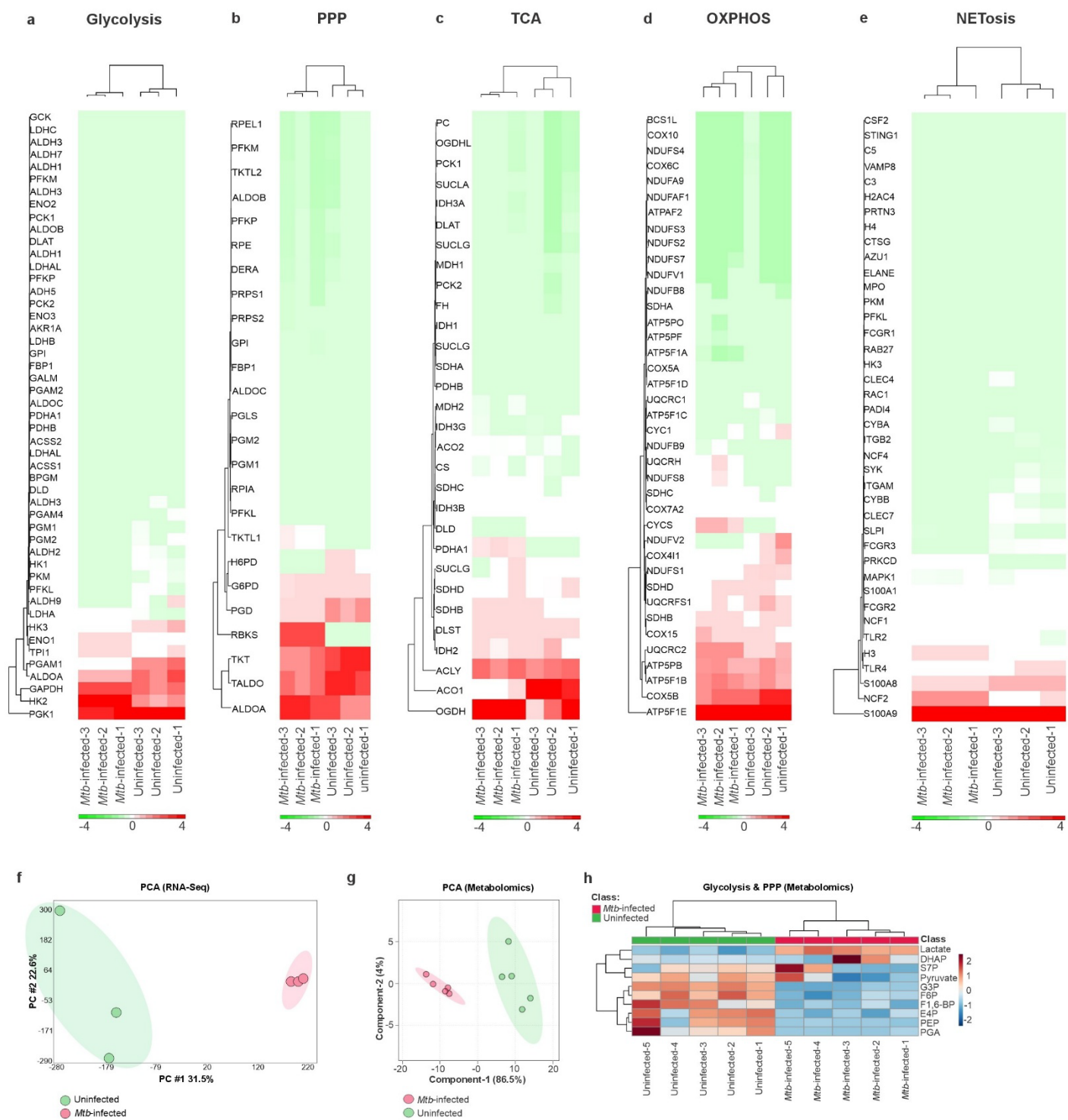

**Extended Data Fig. 9 | Transcriptomic and metabolomic profiling of human neutrophils infected with *Mtb*.**

**a-e**, RNA-Seq analysis of human neutrophils infected with *Mtb* (MOI of 5) for 1 hr. Compared to uninfected controls, gene expression signatures in *Mtb*-infected neutrophils were enriched for glycolysis (**a**), the PPP (**b**), the TCA cycle (**c**), OXPHOS (**d**), and NETosis (**e**). **f**, Principal component analysis (PCA) of total neutrophil transcriptomes shows clear separation between *Mtb*-infected and uninfected groups. **g**, PCA analysis of metabolomic profiles further distinguishes infected from uninfected neutrophils. **h**, cluster enrichment analysis of glycolytic and PPP intermediates in *Mtb*-infected neutrophils. RNA-seq data represents 3 biological replicates; metabolomic profiling represents 5 biological replicates.

| Patient ID | Age | Gender | Disease/Treatment | Resected Lung |
| --- | --- | --- | --- | --- |
| 1 (healthy) | 53 | Male | RA | N/A |
| 2 (healthy) | 54 | Female | Epilepsy | N/A |
| 1 | 44 | Male | TB/DS | Right |
| 4 | 28 | Male | TB/DS | Right |
| 5 | 45 | Male | TB/DS | Left |
| 6 | 34 | Male | TB/MDR | Right |
| 7 | 28 | Female | TB/N/A | Right |
| 8 | 29 | Male | TB/DS | Right |
| 9 | 33 | Female | TB/N/A | Right |
| 10 | 47 | Male | TB/DS | Left |
| 11 | 39 | Female | TB/N/A | Right |
| 12 | 33 | Female | TB/DS | Left |
| 13 | 35 | Female | TB/MDR | Left |
| 14 | 37 | Female | TB/MDR | Left |
| 15 | 33 | Female | TB/DS | Left |
| 16 | 40 | Male | TB/DS | Left |
| 17 | 51 | Male | TB/DS | Right |
| 18 | 32 | Male | TB/N/A | Right |
| 19 | 34 | Female | TB/DS | Right |

**Extended Data Table 1 | Clinical characteristics of human subjects (related to Fig. 1).** Male and female patients were recruited at King DinuZulu Hospital Complex, Durban, South Africa, with written informed consent obtained from all participants. Acronyms: MDR, multi-drug resistant TB; DS, drug-sensitive TB; N/A, information not available.

| Gene Name | Necrotic<br>Granuloma Vs<br>Healthy Control | Log2 Fold Change | p-value |
| --- | --- | --- | --- |
| ACTR2 | Higher Abundance | 0.864241769 | 3.97E-05 |
| ADA2 | Higher Abundance | 1.433766046 | 1.63E-06 |
| ADAM10 | Higher Abundance | 2.381432024 | 2.34E-05 |
| AGRE5 | Higher Abundance | 1.39450124 | 5.42E-07 |
| ALOX5 | Higher Abundance | 2.098981112 | 4.65E-04 |
| AMPD3 | Higher Abundance | 2.553470355 | 2.74E-05 |
| APRT | Higher Abundance | 1.404828881 | 1.27E-09 |
| ARHGAP9 | Higher Abundance | 1.997701482 | 4.65E-06 |
| ARL8A | Higher Abundance | 0.849391375 | 1.16E-05 |
| ARSA | Higher Abundance | 1.163629699 | 1.97E-09 |
| ATP6V0A1 | Higher Abundance | 1.830060307 | 1.12E-06 |
| ATPV1D | Higher Abundance | 0.659751066 | 6.32E-06 |
| AZU1 | Higher Abundance | 1.788902752 | 5.23E-04 |
| ATG7 | Higher Abundance | 0.396000413 | 0.006472184 |
| CALML5 | Higher Abundance | 2.301067424 | 4.60E-07 |
| CAMP | Higher Abundance | 3.293890352 | 8.27E-05 |
| CD14 | Higher Abundance | 1.757204637 | 1.17E-09 |
| CD63 | Higher Abundance | 1.352687682 | 7.55E-06 |
| CDA | Higher Abundance | 1.799025673 | 5.31E-06 |
| CFD | Higher Abundance | 3.785370252 | 3.07E-06 |
| CKAP4 | Higher Abundance | 1.481499587 | 4.24E-10 |
| COPB1 | Higher Abundance | 1.024510372 | 7.19E-09 |
| COTL1 | Higher Abundance | 1.167097006 | 8.20E-10 |
| CPNE3 | Higher Abundance | 1.226641871 | 4.50E-06 |
| CTSG | Higher Abundance | 0.734812642 | 5.10E-05 |
| CTSS | Higher Abundance | 0.894102729 | 3.57E-06 |
| CYBA | Higher Abundance | 2.890676423 | 1.55E-07 |
| CYBB | Higher Abundance | 2.139727538 | 3.83E-12 |
| CASP1 | Higher Abundance | 1.164931036 | 8.70E-08 |
| CEACAM8 | Higher Abundance | 2.353048885 | 0.00159846 |
| CYBA / NOXO2 | Higher Abundance | 2.890676423 | 1.55E-07 |
| CYBB | Higher Abundance | 2.139727538 | 3.83E-12 |
| DDOST | Higher Abundance | 0.971542056 | 1.88E-10 |
| DDX3X | Higher Abundance | 1.261470317 | 7.66E-08 |
| DIAPH1 | Higher Abundance | 0.683390241 | 2.61E-05 |
| DOCK2 | Higher Abundance | 2.2533322 | 1.32E-05 |
| DSG1 | Higher Abundance | 3.736855932 | 1.36E-05 |
| DSP | Higher Abundance | 1.594339788 | 1.81E-04 |
| EEF1A1 | Higher Abundance | 0.977912919 | 7.04E-11 |
| EEF2 | Higher Abundance | 1.452481898 | 4.47E-13 |
| ELANE | Higher Abundance | 2.190575287 | 1.60E-07 |
| ERP44 | Higher Abundance | 0.563253984 | 1.03E-06 |
| FGL2 | Higher Abundance | 0.981556705 | 3.06E-04 |
| FGR | Higher Abundance | 2.260286424 | 9.12E-10 |
| FLG2 | Higher Abundance | 1.498997669 | 3.03E-04 |
| FPR1 | Higher Abundance | 1.463537248 | 9.98E-06 |
| FTH1 | Higher Abundance | 3.295347548 | 1.91E-10 |
| FTL | Higher Abundance | 2.01329027 | 7.11E-09 |
| GCA | Higher Abundance | 1.955267772 | 1.97E-06 |
| GHDC | Higher Abundance | 0.809791909 | 1.20E-04 |
| GM2A | Higher Abundance | 1.528699012 | 1.11E-04 |
| GMFG | Higher Abundance | 1.15982254 | 1.14E-06 |
| GSN | Higher Abundance | 1.219841647 | 1.62E-12 |
| GYG1 | Higher Abundance | 0.998963657 | 4.93E-06 |
| HK3 | Higher Abundance | 3.459436173 | 1.43E-11 |
| HLA-C | Higher Abundance | 1.149503182 | 5.69E-04 |
| HVCN1 | Higher Abundance | 1.958234137 | 1.85E-07 |
| H2AZ1 | Higher Abundance | 0.601337432 | 0.020626627 |
| H4-16 | Higher Abundance | 0.586172756 | 5.93E-04 |
| IMPDH1 | Higher Abundance | 1.946881856 | 1.56E-06 |
| IMPDH2 | Higher Abundance | 1.355488761 | 2.78E-05 |
| IQGAP1 | Higher Abundance | 0.630470881 | 5.14E-08 |
| IQGAP2 | Higher Abundance | 2.436828901 | 7.21E-08 |
| ITGAL | Higher Abundance | 2.31830491 | 3.98E-14 |
| ITGAM | Higher Abundance | 3.366334351 | 5.90E-11 |
| ITGAX | Higher Abundance | 3.800803124 | 1.54E-11 |
| ITGB2 | Higher Abundance | 2.38739884 | 7.76E-11 |
| ICAM1 | Higher Abundance | 0.82057799 | 0.001072313 |
| ITGAM | Higher Abundance | 3.366334351 | 5.90E-11 |
| KRT1 | Higher Abundance | 1.80395946 | 1.70E-06 |
| LAMTOR1 | Higher Abundance | 3.921490883 | 3.38E-06 |
| LAMTOR2 | Higher Abundance | 0.774271316 | 4.15E-04 |
| LCN2 | Higher Abundance | 2.046367021 | 1.54E-04 |
| LILRB2 | Higher Abundance | 1.618183483 | 6.51E-05 |
| LTF | Higher Abundance | 1.699451932 | 9.12E-04 |
| LYZ | Higher Abundance | 2.27522372 | 1.83E-07 |
| MAGT1 | Higher Abundance | 1.84341075 | 4.45E-04 |
| MAN2B1 | Higher Abundance | 1.356931755 | 3.94E-04 |
| MAPK1 | Higher Abundance | 1.324167221 | 1.20E-08 |
| MCEMP1 | Higher Abundance | 0.97964172 | 2.42E-04 |
| MGAM | Higher Abundance | 1.117671906 | 4.56E-05 |
| MPO | Higher Abundance | 2.393040191 | 2.66E-05 |
| MVP | Higher Abundance | 0.517842769 | 5.06E-05 |
| MAPK1 | Higher Abundance | 1.324167221 | 1.20E-08 |
| NCKAP1L | Higher Abundance | 2.568884026 | 1.22E-08 |
| NCF2 | Higher Abundance | 4.140276276 | 7.01E-10 |
| NFKB1 | Higher Abundance | 0.336293832 | 0.002811036 |
| OLFM4 | Higher Abundance | 2.736171173 | 2.79E-06 |
| OSTF1 | Higher Abundance | 0.854071948 | 3.08E-06 |
| PA2G4 | Higher Abundance | 0.87126677 | 2.61E-10 |
| PADI2 | Higher Abundance | 0.607955917 | 3.32E-05 |
| PDXK | Higher Abundance | 1.017588442 | 6.80E-07 |
| PFKL | Higher Abundance | 1.411626536 | 3.65E-10 |
| PKM | Higher Abundance | 2.519262973 | 2.69E-11 |
| PLEKHO2 | Higher Abundance | 1.384234078 | 5.63E-05 |
| PRKCD | Higher Abundance | 1.400451406 | 6.65E-04 |
| PRTN3 | Higher Abundance | 1.369521676 | 1.62E-05 |
| PSMD1 | Higher Abundance | 0.691572552 | 6.64E-06 |
| PTGES2 | Higher Abundance | 0.563409287 | 7.63E-04 |
| PTPN6 | Higher Abundance | 2.685177307 | 2.40E-14 |
| PTPRC | Higher Abundance | 2.520211965 | 1.64E-12 |
| PADI4 | Higher Abundance | 1.789815404 | 0.001636988 |
| PLCG2 | Higher Abundance | 2.618670304 | 6.65E-10 |
| PRKCB | Higher Abundance | -8.585953885 | 7.82E-05 |
| QPCT | Higher Abundance | 0.618107857 | 8.91E-04 |
| QSOX1 | Higher Abundance | 2.408874052 | 8.88E-08 |
| RAB10 | Higher Abundance | 0.726131131 | 4.67E-12 |
| RAB14 | Higher Abundance | 0.259498279 | 9.63E-04 |
| RAB27A | Higher Abundance | 1.600932142 | 1.02E-06 |
| RAB31 | Higher Abundance | 1.252643598 | 3.86E-08 |
| RAB7A | Higher Abundance | 0.917786311 | 4.60E-09 |
| RAC1 | Higher Abundance | 0.917786311 | 4.60E-09 |
| RETN | Higher Abundance | 1.884655148 | 2.59E-04 |
| RHOA | Higher Abundance | 0.607551059 | 4.02E-04 |
| RHOG | Higher Abundance | 2.430486345 | 1.28E-07 |
| RNASE3 | Higher Abundance | 1.401658516 | 2.36E-07 |
| RAC2 | Higher Abundance | 2.349535212 | 2.79E-05 |
| S100A11 | Higher Abundance | 1.138056571 | 1.17E-08 |
| S100A12 | Higher Abundance | 1.382238021 | 4.29E-04 |
| S100P | Higher Abundance | 1.575990938 | 6.82E-05 |
| SDCBP | Higher Abundance | 3.306601345 | 1.82E-06 |
| SERPINB3 | Higher Abundance | 3.856179434 | 2.56E-04 |
| GLUT 3 | Higher Abundance | 3.205142442 | 2.66E-07 |
| SRP14 | Higher Abundance | 0.409997511 | 9.74E-05 |
| STING1 | Higher Abundance | 1.371522376 | 2.74E-06 |
| STOM | Higher Abundance | 1.291830887 | 7.48E-06 |
| TMEM30A | Higher Abundance | 1.222732698 | 6.35E-04 |
| TOM1 | Higher Abundance | 1.017815162 | 3.03E-04 |
| TRAPPC1 | Higher Abundance | 1.414762519 | 6.98E-10 |
| TTR | Higher Abundance | 1.262925741 | 5.06E-06 |
| TUBB | Higher Abundance | 0.52142565 | 1.47E-06 |
| TXNDC5 | Higher Abundance | 1.085993495 | 2.11E-07 |
| UNC13D | Higher Abundance | 2.28448256 | 7.62E-08 |
| VAT1 | Higher Abundance | 0.480373967 | 9.45E-06 |
| VPS35L | Higher Abundance | 1.457745553 | 9.28E-07 |
| AKT1 | Lower Abundance | -1.04031615 | 0.008215029 |
| DPP7 | Lower Abundance | -0.926536041 | 4.05E-04 |
| DYNLL1 | Lower Abundance | -1.451666145 | 3.80E-05 |
| FUCA1 | Lower Abundance | -0.919531079 | 9.65E-04 |
| HMGB1 | Lower Abundance | -1.061148853 | 8.27E-10 |
| HP | Lower Abundance | -3.107004634 | 5.92E-11 |
| HSPA1A | Lower Abundance | -1.029769867 | 4.45E-05 |
| HDAC1 | Lower Abundance | -0.766266869 | 0.005774772 |
| HMGB1 | Lower Abundance | -1.061148853 | 8.27E-10 |
| LAMTOR3 | Lower Abundance | -0.531780156 | 1.04E-04 |
| LRG1 | Lower Abundance | -2.424772347 | 1.64E-06 |
| MLEC | Lower Abundance | -2.086120557 | 7.75E-04 |
| MME | Lower Abundance | -0.953368356 | 0.052504483 |
| NEU1 | Lower Abundance | -2.279660931 | 1.94E-05 |
| ORM1 | Lower Abundance | -1.788281385 | 1.24E-13 |
| OSCAR | Lower Abundance | -1.710512508 | 4.69E-04 |
| PGM2 | Lower Abundance | -0.900067527 | 4.13E-05 |
| PRDX6 | Lower Abundance | -2.106497627 | 5.00E-04 |
| PSMA5 | Lower Abundance | -0.758424115 | 1.93E-06 |
| PSMB1 | Lower Abundance | -1.011458989 | 5.67E-04 |
| SERPINA1 | Lower Abundance | -1.660318772 | 8.87E-04 |
| SERPINA3 | Lower Abundance | -1.33069937 | 2.08E-07 |
| SERPINB6 | Lower Abundance | -1.413146211 | 1.59E-04 |
| SIGLEC9 | Lower Abundance | -1.239956758 | 9.49E-04 |
| SLPI | Lower Abundance | -1.131178306 | 5.00E-08 |
| SPTAN1 | Lower Abundance | -2.210790024 | 4.73E-04 |
| SELPLG | Lower Abundance | -0.682242496 | 0.015158711 |
| SOD2 | Lower Abundance | -0.785443998 | 0.004679355 |
| TUBB4B | Lower Abundance | -0.802451875 | 6.69E-04 |
| TLR2 | Lower Abundance | -0.39918178 | 0.021537863 |
| ALAD | Lower Abundance | -2.825954183 | 9.36E-08 |
| APEH | Lower Abundance | -1.65276325 | 5.52E-04 |
| ARG1 | Lower Abundance | -4.965963843 | 0.013165928 |
| C3 | Lower Abundance | -1.568343784 | 5.70E-06 |
| CAT | Lower Abundance | -2.752078878 | 8.93E-04 |
| CD59 | Lower Abundance | -1.761151909 | 5.96E-04 |
| CD93 | Lower Abundance | -2.199101976 | 4.46E-20 |
| CTSC | Lower Abundance | -0.324406558 | 6.28E-04 |

### Extended Data Table 2 | Neutrophil-associated proteins in necrotic human TB granulomas

(related to Fig. 1). Proteomic analysis of laser-capture micro-dissected necrotic TB granulomas identified 1,227 differentially abundant proteins compared with non-TB control lung tissue. Among these, 173 were neutrophil-associated proteins, of which 135 were present at higher abundance in necrotic TB lesions and 38 at lower abundance relative to controls. The table lists protein gene names, relative abundance classification (high or low), $\log_2$ fold change (necrotic TB versus non-TB lung), and corresponding $p$ values.

| Patient ID | Age | Gender | Patient Status | Resected Lung |
| --- | --- | --- | --- | --- |
| Control-1 | 54 | Female | Cancer | Right |
| Control-2 | 68 | Male | Cancer | Left |
| Control-3 | 63 | Female | Cancer | Left |
| Control-4 | 66 | Male | Cancer | Left |
| Control-5 | 76 | Female | Cancer | Right |
| Control-6 | 58 | Female | Cancer | Left |
| Control-7 | 56 | Female | N/A | Right |
| Control-8 | 31 | Male | Cancer | Right |
| Control-9 | 56 | Male | Cancer | Right |
| Control-10 | 65 | Female | Cancer | Left |
| Control-11 | 50 | Male | Cancer | Left |
| TB-1 | 35 | Male | MDR | Left |
| TB-2 | 37 | Female | DS | Right |
| TB-3 | 18 | Male | DS | Right |
| TB-4 | 35 | Female | DS | Left |
| TB-5 | N/A | N/A | N/A | Left |
| TB-6 | 51 | Male | DS | Right |
| TB-7 | 21 | Male | MDR | Right |
| TB-8 | 42 | Female | MDR | Left |
| TB-9 | 31 | Male | DS | Right |
| TB-10 | 41 | Male | DS | Left |
| TB-11 | 33 | Female | DS | Right |
| TB-12 | 35 | Male | N/A | Left |
| TB-13 | 24 | Female | XDR | Left |
| TB-14 | 30 | Female | DS | Left |
| TB-15 | 33 | Female | MDR | Right |
| TB-16 | 24 | Female | DS | Left |
| TB-17 | 53 | Male | XDR | Right |
| TB-18 | 30 | Female | DS | Right |
| TB-19 | 47 | Male | DS | Right |

**Extended Data Table 3 | Clinical characteristics of human subjects** (related to Extended Data Fig.

1). Male and female patients were recruited at King DinuZulu Hospital Complex, Durban, South Africa, with written informed consent obtained from all participants. Acronyms: MDR, multi-drug resistant TB; DS, drug-sensitive TB; N/A, information not available.
